# Comprehensive characterisation of IAA inactivation pathways reveals the impact of glycosylation on auxin metabolism and plant development

**DOI:** 10.64898/2026.01.19.700167

**Authors:** Rubén Casanova-Sáez, Aleš Pěnčík, Federica Brunoni, Anita Ament, Pavel Hladík, Asta Žukauskaitė, Jan Šimura, Ute Voß, Ondřej Novák, Malcolm Bennett, Karin Ljung, Eduardo Mateo-Bonmatí

**Affiliations:** Umeå Plant Science Centre, Department of Forest Genetics and Plant Physiology, Swedish University of Agricultural Sciences, SE-901 83 Umeå, Sweden; Laboratory of Growth Regulators, Faculty of Science, Palacký University, Šlechtitelů 27, Olomouc CZ-77900, Czech Republic; Laboratory of Growth Regulators, Institute of Experimental Botany, The Czech Academy of Sciences, Šlechtitelů 27, Olomouc CZ-77900, Czech Republic; Department of Chemical Biology, Faculty of Science, Palacký University, Šlechtitelů 27, CZ-77900 Olomouc, Czech Republic; Division of Plant and Crop Sciences, School of Biosciences, University of Nottingham, Sutton Bonington Campus, Loughborough, LE12 5RD, UK; Centro de Biotecnología y Genómica de Plantas (CBGP), Universidad Politécnica de Madrid (UPM), Instituto Nacional de Investigación y Tecnología Agraria y Alimentaria (INIA)/CSIC, Pozuelo de Alarcón, Madrid 28223, Spain

**Keywords:** Arabidopsis, IAA metabolism, DAO1, GH3, UGTs, FLC, RNA-seq

## Abstract

Together with biosynthesis and transport, inactivation regulates the concentration of indole-3-acetic acid (IAA), a key auxinic compound with a myriad of functions in plant development. Main inactive IAA metabolites are categorised into oxidised forms and ester- or amide-linked conjugates. DIOXYGENASE FOR AUXIN OXIDATION1 (DAO1) and DAO2, 2-oxoglutarate and iron-dependent dioxygenases, contribute to IAA oxidative inactivation in collaboration with group II GRETCHEN HAGEN3 (GH3) IAA-amido synthetases, while a group of UDP-glycosyltransferases (UGTs) conjugate IAA to sugars. To study the IAA inactivation routes, we generated combinatorial mutants between all group II *GH3*s (*gh3oct*) and *DAO1* or *DAO2*, as well as between the *DAOs* and main *UGT*s. *In vivo* [^13^C_6_]IAA feeding experiments traced the metabolic fate of the exogenously applied IAA, supporting the main IAA inactivation pathway, in which DAO acts downstream of GH3s. They also indicated that UGT-mediated IAA glycosylation is more important than previously assumed for modulating IAA levels and plant development. Our metabolic and transcriptomic data further revealed that *gh3oct* may still produce some *GH3* activity, explaining previous reported phenotypic inconsistencies. Our data additionally suggest that other not yet identified metabolic activities might play a role in IAA overproducing plants, and that the premature downregulation of flowering time integrators like *FLOWERING LOCUS C* (*FLC*) likely underlies the early flowering of *gh3oct* and *gh3oct dao1* plants.

## INTRODUCTION

As biochemical messengers, hormones synchronize developmental stages with environmental conditions to ensure the survival and reproduction of eukaryotic organisms. This is particularly important in the plant lineage since their sessile nature hampers other possible responses. Among plant hormones, indole-3-acetic acid (IAA), the most important form of auxin, has been found to play a role in many developmental transitions^1,2^ as well as in the integration of external signals^3,4^ and stress responses^5,6^. Auxińs action depends on its cellular concentration. Contrary to the transport-centric original dogma^7,8^, it is now believed that local auxin concentration results from concerted action of at least three convergent and intertwined pathways: auxin (polar) transport, auxin biosynthesis, and auxin inactivation^1,9^. All of them are complex pathways with multiple genes and cross-regulatory relationships.

IAA inactivation involves modifications of its molecular structure, rendering it unrecognizable to the signalling and transport machinery. Several chemical modifications have been observed on the IAA molecule. IAA can be inactivated by methylation (MeIAA)^10^ and via ester-linked and amide-linked conjugation^1^. The most abundant inactive form is the ester-linked IAA-glucose (IAA-glc), present at high levels in different plants, especially in seedlings and seeds^11-13^. Enzymes catalysing the glycosylation of IAA belong to the family of UDP-glycosyltransferases. Despite being a large family, only three UGTs have been identified as playing a role in IAA inactivation in Arabidopsis: UGT84B1, UGT74D1, and UGT76E5^14-16^. Amide-linked conjugates comprise a group of molecules where IAA is conjugated mainly to an amino acid (IAA-aa) or to small peptides and proteins. The formation of IAA-aa conjugates is catalysed by the GRETCHEN HAGEN3 (GH3) family of acyl acid amido synthetases^17^, another big gene family containing 19 members in Arabidopsis, clustered in three functional groups^18^. Group II GH3 members are known to catalyse the formation of the different IAA-aa conjugates, which include IAA-leucine, IAA-alanine, IAA-phenylalanine, IAA-aspartate (IAA-Asp), or IAA-glutamate (IAA-Glu)^19^. Oxidized counterparts of these IAA-aa conjugates, such as 2-oxindole-3-acetic acid-aspartate (oxIAA-Asp) and oxIAA-Glu are also formed^20,21^.

Another inactive IAA is found in the oxidized form (oxIAA), initially reported to be generated by the activity of the 2-oxoglutarate and iron-dependent dioxygenases (2OGD) DIOXYGENASE FOR AUXIN OXIDATION 1 (DAO1) and DAO2^22-24^. Recent reports, however, indicate that DAOs may operate mainly downstream of GH3s by converting IAA-aa conjugates into oxIAA-aa conjugates^20,25^, later transformed into oxIAA by the action of amidohydrolases like IAA-LEUCINE RESISTANT1 (ILR1)^25^.

To elucidate the role of the main auxin inactivation routes (Figure 1) in regulating IAA levels and plant development, we generated a comprehensive set of genetic mutants. We examined phenotypic outcomes and quantified the metabolic fate of [^13^C_6_]-labelled IAA across all genotypes to trace the activity of distinct inactivation routes and their associated developmental implications. We also conducted transcriptomic analyses of the IAA response in two genotypes impaired in the primary inactivation mechanisms. Our results outline an important role of the IAA glycosylation pathway in IAA metabolism and plant development, and support the notion that GH3s and DAOs act in a primary and consecutive pathway for IAA inactivation. Finally, our findings suggest the existence of additional, yet unidentified, players contributing to IAA inactivation.

**Figure 1.**
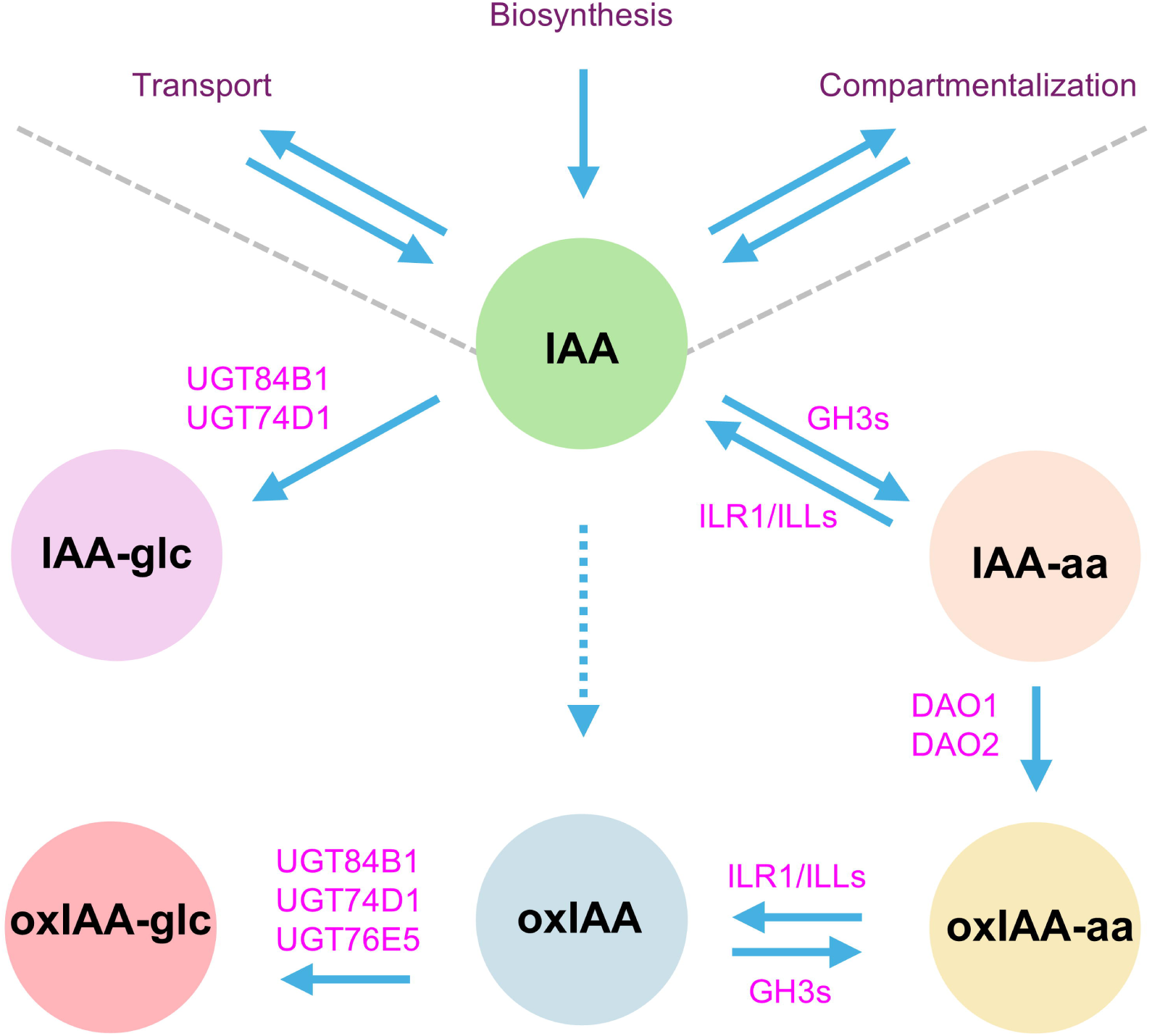
Main pathways for indole-3-acetic acid (IAA) metabolic inactivation in Arabidopsis. Along with biosynthesis, transport and subcellular compartmentalization, inactivation defines the cellular levels of IAA. IAA metabolites are indicated in circular boxes. Enzymes catalyzing the metabolic reactions are indicated in magenta. IAA-glc: IAA-glucose (indole-3-acetyl-β-D-glucopyranoside); IAA-aa: IAA-amino acid conjugates, such as IAA-Asp (indole-3-acetyl-L-aspartic acid) and IAA-Glu (indole-3-acetyl-L-glutamic acid); oxIAA-aa: conjugates of oxIAA (2-oxoindole-3-acetic acid) with amino acids; oxIAA-glc: oxIAA-glucose (2-oxoindole-3-acetyl-β-D-glucopyranoside). GH3s: GRETCHEN HAGEN3 IAA acyl acid amido synthetases. DAO1/2: DIOXYGENASE FOR AUXIN OXIDATION 1/2. ILR1/ILLs: IAA-LEUCINE RESISTANT1/ILR1-LIKE enzymes. UGT84B1/74D1/76E5: URIDINE-DIPHOSPHATE GLYCOSYLTRANSFERASE 84B1/74D/76E5.

## RESULTS

### Generation of a genetic toolkit to study IAA inactivation

To investigate the contribution of the major IAA inactivation pathways (Figure 1) to plant development, we undertook a reverse genetic approach to generate mutants that disrupt these pathways individually and in combination. For this purpose, we combined previously generated lines with newly generated CRISPR/Cas9-induced mutants. Previously, we identified *UGT84B1* and *UGT74D1* as the main UDP-glycosyltransferases responsible for conjugating glucose to both IAA and oxIAA^14^. To effectively disrupt the IAA glycosylation pathway, we generated a *ugt84b1 ugt74d1* double mutant by CRISPR/Cas9-mediated knock-out of *UGT84B1*^14^ in the T-DNA *ugt74d1* mutant background^15^.

In parallel, we generated a *dao1 dao2* double mutant by introducing a CRISPR/Cas9-mediated deletion of *DAO1* in the *dao2-1* insertion allele background^23^. This deletion (hereafter referred to as *dao1-4*) removed a 712 bp genomic fragment of within *DAO1*, resulting in a frameshift change at the amino acid 75 and a premature stop codon at the amino acid 87. This truncation effectively eliminates the 2-oxoglutarate and iron-dependent dioxygenase (2OGD) domain, essential for enzymatic function, from the resulting mutant protein (Figure 2).

**Figure 2.**
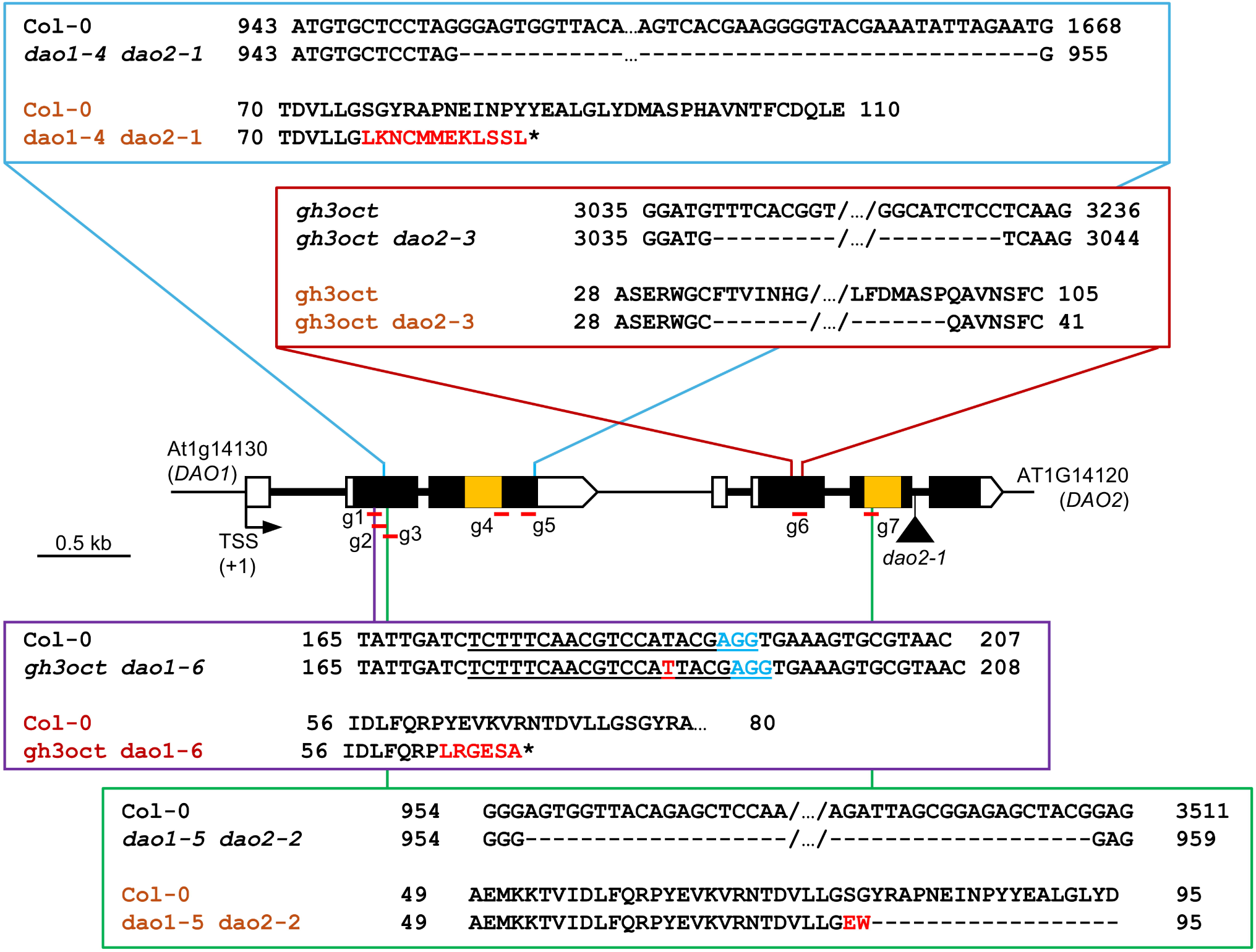
Schematic representation of the multiple mutants generated. Illustration of the editing events isolated after targeting the CRISPR/Cas9 system to the genomic region encompassing the *DAO1* and *DAO2* genes. Architecture of the *DAO1* and *DAO2* genes, and illustration of the nature and location of the mutations studied in this work. Black boxes indicate exons, and lines indicate introns. White boxes represent untranslated regions. The triangle represents a T-DNA insertion (*dao2-1*). The orange stripes represent the region encoding for the oxoglutarate/iron-dependent dioxygenase domain (2OGD; IPR005123). Red horizontal bars represent sgRNAs (g1-to-g7; not drawn to scale) used to edit the region. The blue square magnifies the *DAO1* deletion found in the *dao2-1* genetic background. Genotypes written in black refer to genomic regions, while genotypes in brown refer to the proteins. The purple square explains the single-nucleotide variant, which causes a premature stop codon found in the *gh3oct* background. Highlighted in light blue is the NGG motif of the sgRNA. The green square shows the large genomic deletion encompassing *DAO1* and *DAO2* in the *ugt74d1* background. Red letters represent either nucleotides or amino acids absent in the wild type. Asterisks represent a premature stop codon. Numbers close to sequences indicate the position of the nucleotide from the transcriptional start site (TSS) of *DAO1* or the amino acid position. Scale bar indicates 0.5 kb.

To simultaneously disrupt the IAA glycosylating and oxidative pathways, we used CRISPR/Cas9 to induce a 2.5-kb genomic deletion encompassing both *DAO1* and *DAO2* genes in the *ugt74d1* background. This deletion removed the 2OGD domains from both *DAO1* and *DAO2* (Figure 2; *dao1-5 dao2-2*). The resulting deletion was then combined with the *ugt84b1 ugt74d1* double mutant by crossing, yielding the triple mutant *ugt84b1 dao1-5 dao2-2*, as well as the quadruple mutant *dao1-5 dao2-2 ugt84b1 ugt74d1*.

To examine the interaction between the conjugation and oxidation pathways for IAA inactivation, we further disrupted either *DAO1* or *DAO2* in the *gh3.1 gh3.2 gh3.3 gh3.4 gh3.5 gh3.6 gh3.9 gh3.17* octuple mutant (*gh3oct*)^19^, generating the *gh3oct dao1-6* and *gh3oct dao2-3* nonuple mutants. The isolated mutation in *DAO1* consisted of a single-nucleotide insertion at position 187 of the *DAO1* transcriptional unit, resulting in a frameshift and a premature stop codon after residue 68 (Figure 2). The *dao2-3* allele was generated via CRISPR/Cas9-mediated deletion of 192 nucleotides, resulting in the loss of 64 amino acids from the DAO2 protein (Figure 2). Together, these lines constitute a comprehensive set of combinatorial mutants to dissect the roles of the IAA inactivation pathways in auxin metabolism and plant development.

### Phenotypic analyses reveal unique contributions of IAA glycosylases and DAO2 to plant development

We then carried out phenotypic analyses of the complete set of generated mutant lines. Consistent with previous reports, *dao1-1* and *dao2-1* single mutants exhibited virtually no root phenotype or hypocotyl phenotype (Figure 3a-c)^22,23^. In contrast, *dao1-4 dao2-1* double mutants displayed a mild, high auxin-associated phenotype of increased primary root length (PRL; Figure 3a; Figure S1). This reveals DAO1 and DAO2 having at least partially overlapping functions and being able to compensate for each other’s loss, despite DAO1 being the predominant oxidase in root IAA metabolism^22^.

**Figure 3.**
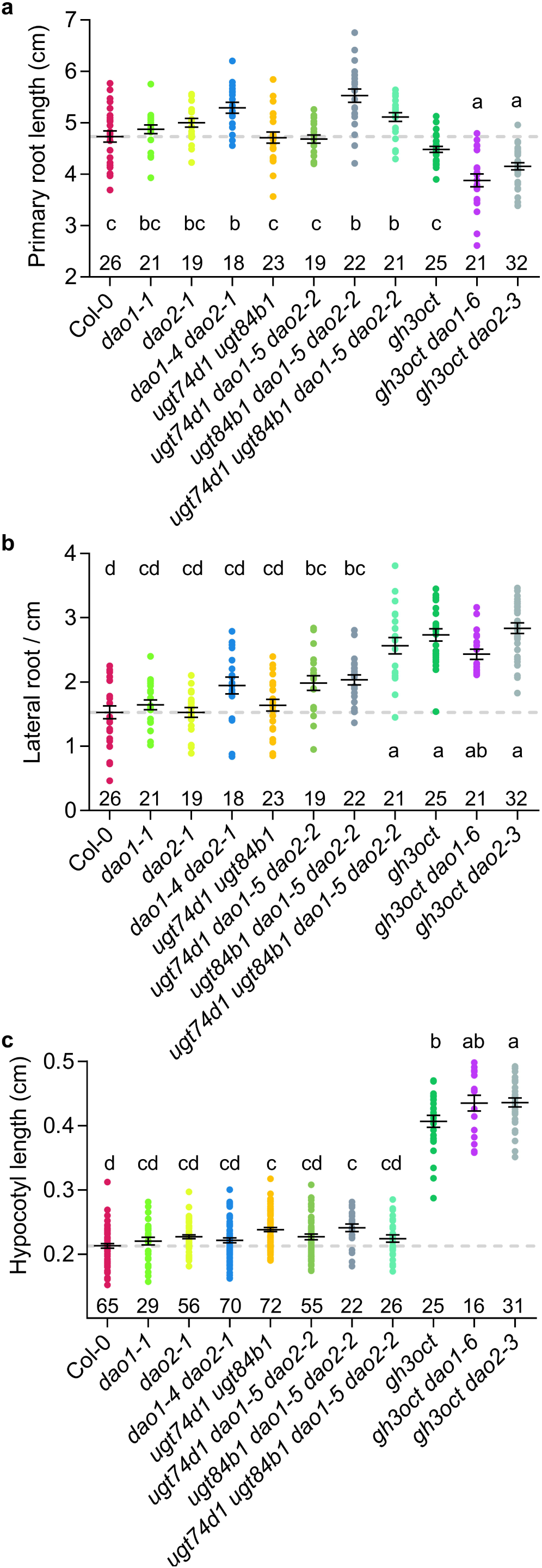
Morphometric traits of the IAA metabolism mutants. (a-c) Different phenotypic variables measured in the assorted genotypes. Dots represent individual scoring of (a) primary root length, (b) lateral root density, and (c) hypocotyl length. (a) and (c) were measured 7 days after stratification (das). (b) was measured 10 das. The average ± standard error of the mean is shown. The grey dashed lines mark the wild-type average. A one-way ANOVA was performed, followed by pairwise comparisons, using Tukeýs HSD test. Conditions marked with different letters (a, b, c) are statistically different (p < 0.05). The number of plants analysed per genotype is shown above the name.

While single *ugt* mutants have previously been reported to lack apparent root phenotypic alterations^14^, *ugt84b1 ugt74d1* double mutant seedlings exhibited increased hypocotyl length (Figure 3c). This indicates that IAA glycosylation, redundantly mediated by UGT84B1 and UGT74D1, plays a significant role in modulating auxin levels in the hypocotyl during seedling development. Remarkably, the lateral root density (LRD) in the quadruple mutant *dao1-5 dao2-2 ugt74d1 ugt84b1* mutant was synergistically increased when compared to *dao1 dao2*, and *ugt74d1 ugt84b1* double mutants (Figure 3b), which suggests that IAA oxidation and glycosylation pathways redundantly contribute to auxin homeostasis and root architecture, while being individually dispensable under standard growth conditions.

In line with the literature, knocking out the *GH3* pathway had more severe developmental impacts compared to *dao* or *ugt* double mutants. The *gh3oct* mutant showed enhanced hypocotyl length and LRD without penalty in the primary root length (Figure 3a-c). Notably, the *gh3oct dao1-6* and *gh3oct dao2-3* nonuple mutants displayed a further reduction in PRL and a greater increase in hypocotyl length compared to *gh3oct* alone (Figure 3b,c), which is indicative of higher IAA levels in nonuple mutant plants and again supports an overlapping yet non-fully redundant role of DAO2 in plant development.

### IAA-glycosylation deficient mutants show significant re-wiring of IAA inactivation routes

To precisely dissect the contributions of the different enzyme sets to IAA metabolism, we quantified the levels of ^13^C_6_-isotope-labelled IAA metabolites, including [^13^C_6_]IAA, [^13^C_6_]IAA-glc, [^13^C_6_]IAA-Glu, [^13^C_6_]IAA-Asp, [^13^C_6_]oxIAA, [^13^C_6_]oxIAA-glc, [^13^C_6_]oxIAA-Glu, and [^13^C_6_]oxIAA-Asp in 7-day-old seedlings from all generated mutant lines after feeding with [^13^C_6_]IAA for 3, 12, and 24 hours (Figure 4 and S2). In parallel, we also measured the steady-state levels of these metabolites in root tissues across genotypes (Figure S3).

**Figure 4.**
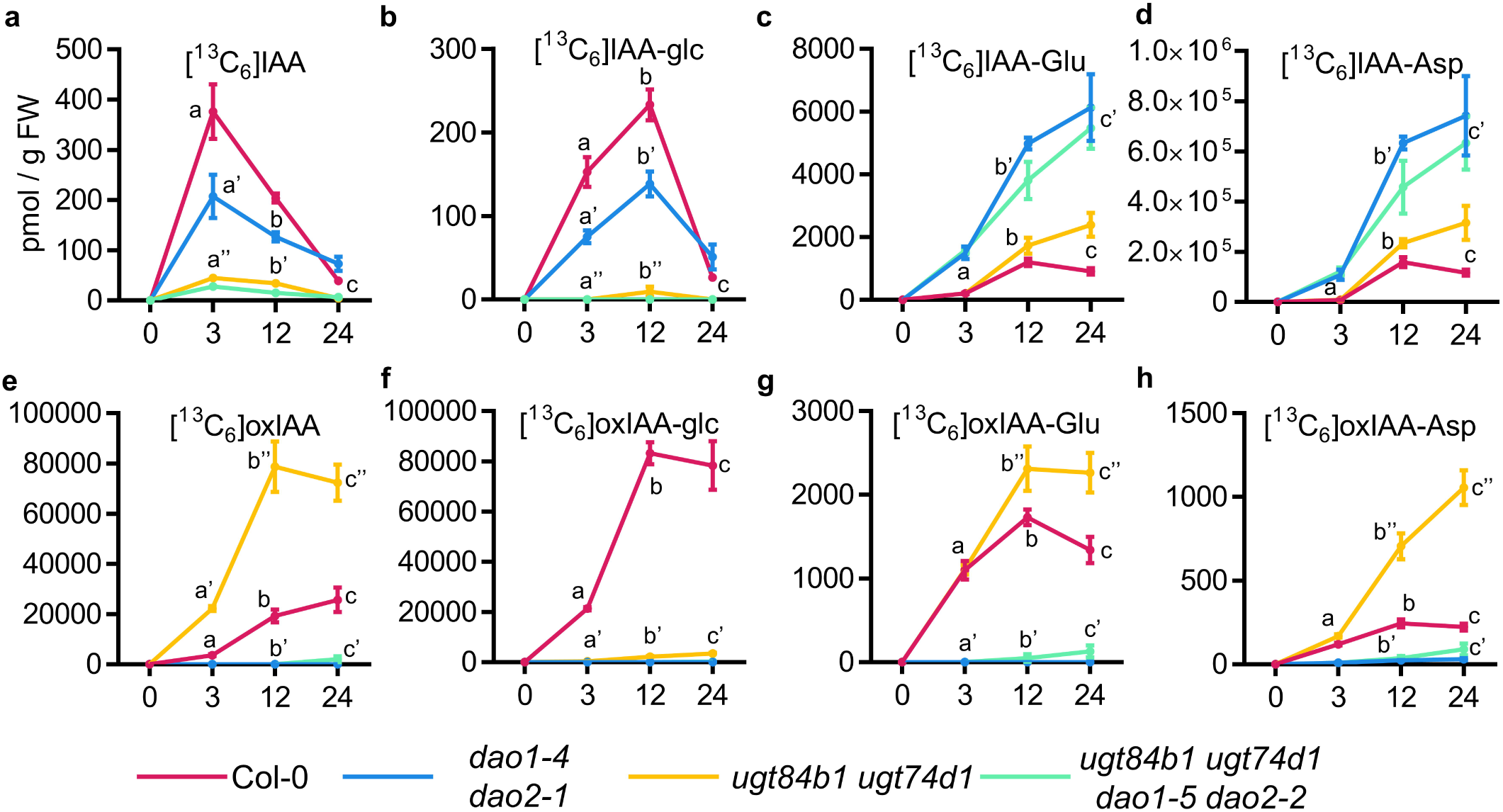
*De novo* synthesis levels of indole-3-acetic acid (IAA) metabolites in different Arabidopsis *DAO* and *UGT* multiple mutant lines. (a-h) Formation of [^13^C_6_]-labelled IAA metabolites in 7-day-old seedlings of the indicated genotypes after incubation with 1 µM [^13^C_6_]IAA for 0, 3, 12, and 24 hours. Dots indicate the mean ± standard error of the mean. Levels in picomoles per gram of fresh weight of (a) [^13^C_6_]IAA, (b) [^13^C_6_]IAA-glc, (c) [^13^C_6_]IAA-Glu, (d) [^13^C_6_]IAA-Asp, (e) [^13^C_6_]oxIAA, (f) [^13^C_6_]oxIAA-glc, (g) [^13^C_6_]oxIAA-Glu, and (h) [^13^C_6_]oxIAA-Asp. For each time point, differences were evaluated by a one-way ANOVA followed by pairwise comparisons using Tukeýs HSD test.

Single *dao1-1* mutant reproduced previously reported metabolic profiles, with a massive accumulation of IAA-aa conjugates^22,23^ and impaired formation of the oxidised forms [^13^C_6_]oxIAA, [^13^C_6_]oxIAA-glc, [^13^C_6_]oxIAA-Glu or [^13^C_6_]oxIAA-Asp (Figure S2). In contrast, the IAA metabolic profile of *dao2-1* closely resembled that of the wild type (Figure S2). Both *de novo* formation of [^13^C_6_]IAA-derived inactive metabolites and the steady-state levels of endogenous IAA metabolites in seedling roots from the *dao1-4 dao2-1* double mutant were comparable to those observed in *dao1-1* (Figures 4, S2, and S3). Interestingly, despite being reduced relative to Col-0, steady-state oxIAA, oxIAA-glc, and oxIAA-aa were still present in both single *dao1-1* and double *dao1-4 dao2-1* mutant plants (Figure S3e-g). One explanation for this phenomenon could be that *DAO2* compensates the lack of *DAO1* in both the *dao1-1* single and the *dao1-1 dao2-1* double, since it was reported that *dao2-1* is a leaky allele of *DAO2*^23^. However, our phenotypic analyses rather disfavour this possibility: other, more deleterious *DAO2* alleles of were obtained in combination with *ugt* mutants without any enhancement of the effects on root growth (Figure 3). Moreover, these additional *dao2* alleles generated in this work show the same feeding (Figure 4) and steady-state levels (Figure S3) of oxIAA, IAA-Glu, IAA-Asp, or oxIAA-glc, suggesting the involvement of non-catalytic oxidation processes and/or the existence of additional, yet unidentified, oxidase activities contributing to IAA turnover.

Not unexpectedly, the formation of [^13^C_6_]IAA-glc was strongly impaired in the *ugt74d1 ugt84b1* mutant (Figure 4b). The statistically significant increase in the formation of other inactive metabolites [^13^C_6_]oxIAA, [^13^C_6_]IAA-Glu, and [^13^C_6_]IAA-Asp (Figure 4c-e) suggests a re-wiring of accumulating IAA towards the oxidative and GH3 pathway as a consequence of disrupting IAA glycosylation. While blocking both the IAA oxidative and glycosylation pathways in the quadruple *ugt84b1 ugt74d1 dao1 dao2* mutant had a stronger impact on lateral root density compared to doubles and triples (Figure 3b), the IAA inactivation network and endogenous levels of inactive IAA forms were similar between the quadruple *ugt84b1 ugt74d1 dao1 dao2* and the double *ugt84b1 ugt74d1* or *dao1 dao2* mutants (Figures 4 and S3). Notably, our data showed identical reduction in [^13^C_6_]IAA accumulation and endogenous IAA levels in both the double *ugt74d1 ugt84b1* and the quadruple *ugt84b1 ugt74d1 dao1 dao2* (Figure 4A and S2A), seemingly resulting from the loss of UGT74D1 (Figures S2A and S3A). This indicates that an impaired IAA glycosylation pathway triggers enhanced metabolic removal of IAA, likely via the GH3 pathway (Figure 4c, d, g, h) or by additional IAA inactivation routes yet to be discovered.

The triple mutant *ugt74d1 dao1 dao2* showed another interesting IAA metabolic rewiring. While, as expected, it showed reduced steady-state levels of oxIAA, lack of oxIAA-glc, and a substantial accumulation of IAA-Glu and IAA-Asp (Figure S3), the feeding experiment revealed a surprising reactivation of the *de novo* formation of both oxIAA and oxIAA-aa (Figure S2e, g, h), with the double *dao1 dao2* mutant unable to produce these metabolites under our experimental conditions (Figure 4e, g, h). Consistently, the accumulation of oxIAA-Asp in the roots of *ugt74d1 dao1 dao2* seedlings was markedly higher than in wild-type or *dao1 dao2* roots (Figure S2g). Likewise, oxIAA-Glu was detectable in the roots of the *ugt74d1 dao1 dao2* but not in those of the *dao1 dao2* mutant (Figure S3f). This further suggests the possible involvement of additional IAA oxidase activities, and points to a complexity and flexibility of the IAA inactivation network higher than currently understood.

### Disruption of the GH3s and DAOs enhances flux through the IAA glycosylation pathway

We next examined the IAA inactivation dynamics in the absence of both the IAA-amino acid conjugation and oxidative routes by performing [^13^C_6_]IAA feeding experiments in the wild type (same profile as above), *gh3oct*, *gh3oct dao1-6,* and *gh3oct dao2-3* genotypes (Figure 5). All three mutant backgrounds exhibited a reduced capacity to metabolize excess IAA, with the effect being more pronounced in *gh3oct dao1-6* plants (Figure 5a). Concurrently, [^13^C_6_]IAA-glc formation was elevated in all three genotypes (Figure 5b). As expected, *gh3oct* plants were severely impaired in redirecting IAA towards IAA-Glu or IAA-Asp (Figure 5c, d), and consistent with DAOs acting downstream of GH3s^20,25^, the formation of oxIAA and oxIAA-glc was strongly reduced in the *gh3oct* background (Figure 5e, f). However, the differential formation of [^13^C_6_]oxIAA-aa conjugates in *gh3oct* and *gh3oct dao1-6* plants (Figure 5g, h), along with the strikingly increased accumulation of [^13^C_6_]IAA-Glu in the *gh3oct dao1-6* mutant (Figure 5c) and the presence of endogenous levels of IAA-Glu in *gh3oct* roots (Figure S3b), suggests that either non-group II GH3s are capable of conjugating IAA under high-hormone conditions, or that a truncated but partially functional GH3 isoform is still produced from one or more of the disrupted loci in the insertional *gh3oct* mutant.

**Figure 5.**
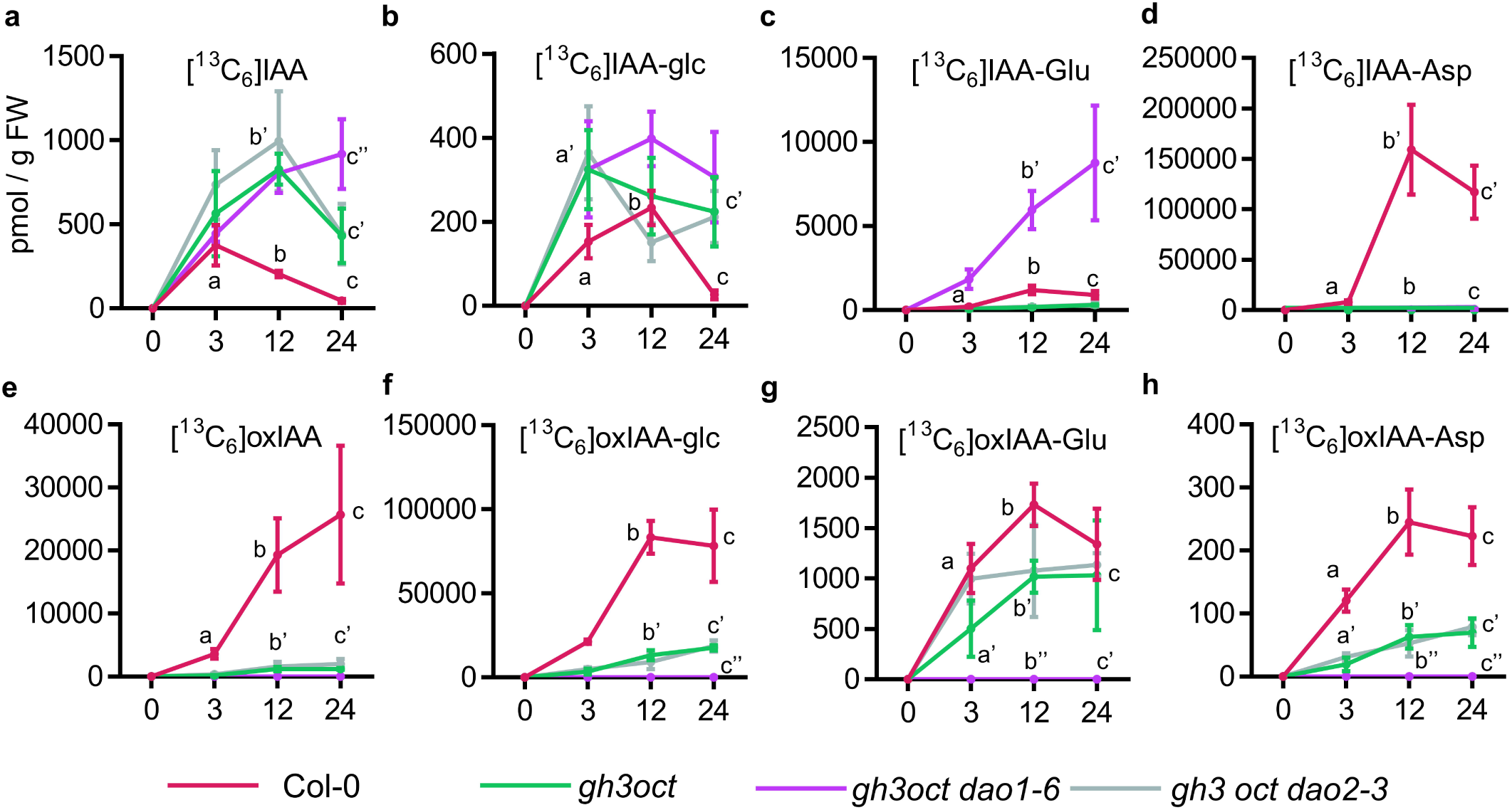
*De novo* synthesis levels of indole-3-acetic acid (IAA) metabolites in different Arabidopsis *GH3* and *DAO* multiple mutant lines. (a-h) Formation of [^13^C_6_]-labelled IAA metabolites in 7-day-old seedlings of the indicated genotypes after incubation with 1 µM [^13^C_6_]IAA for 0, 3, 12, and 24 hours. Dots indicate the mean ± standard error of the mean. (A-H) Levels in picomoles per gram of fresh weight of (a) [^13^C_6_]IAA, (b) [^13^C_6_]IAA-glc, (c) [^13^C_6_]IAA-Glu, (d) [^13^C_6_]IAA-Asp, (e) [^13^C_6_]oxIAA, (f) [^13^C_6_]oxIAA-glc, (g) [^13^C_6_]oxIAA-Glu, and (h) [^13^C_6_]oxIAA-Asp. For each time point, differences were evaluated by a one-way ANOVA followed by pairwise comparisons using Tukeýs HSD test.

To better understand this phenomenon, we assayed the response of *gh3oct* plants to kakeimide (KKI)^26^, a chemical inhibitor of the GH3 activity. We vertically grew Col-0 and *gh3oct* plants for one week in half-strength MS (mock) and in media supplemented with 5 µM of KKI. We then scored the primary root length and lateral root density. As previously reported^19^ and further reproduced in this work (Figure 3a, b), *gh3oct* seedlings exhibited more lateral roots without a reduction in primary root length at 7 days (Figure S4). While one would expect the *gh3oct* mutant to be fully insensitive to KKI treatment, we found that primary root growth in *gh3oct* is inhibited by KKI to a similar extent as in the wild type. Similarly, root branching was also enhanced upon treatment with KKI. These results suggest that residual GH3 conjugation activity remains in the *gh3oct* mutant and can be fully inhibited by KKI. These new observations help resolve the apparent discrepancy between the phenotypes of the *gh3oct* mutant and those of a similar mutant generated by CRISPR/Cas9 and reported at a similar time^27^.

To study the response of the IAA inactivation network to a fully blocked GH3-DAO pathway (Figure 1), we performed a [^13^C_6_]IAA feeding experiments in triple amidohydrolase *ilr1-1 iar3-2 ill2-1* mutant plants in which the IAA-conjugating GH3 activity was chemically inhibited with kakeimide (KKI^26^; Figure 6). Under these conditions, the formation of [^13^C_6_]IAA-aa conjugates was abolished in both wild-type and *ilr1-1 iar3-2 ill2-1* plants (Figure 6c, d). While [^13^C_6_]oxIAA-aa conjugates were only marginally detected in the wild-type, they significantly accumulated in the *ilr1-1 iar3-2 ill2-1* mutant (Figure 6g, h). The increased levels of [^13^C_6_]IAA-glc in GH3-inhibited *ilr1-1 iar3-2 ill2-1* plants indicate a redirection of excess IAA towards the IAA glycosylation pathway (Figure 6b). Remarkably, no [^13^C_6_]oxIAA and [^13^C_6_]oxIAA-glc were produced in *ilr1-1 iar3-2 ill2-1* plants (Figure 6e, f), supporting that DAOs act exclusively in the GH3s pathway^25^.

**Figure 6.**
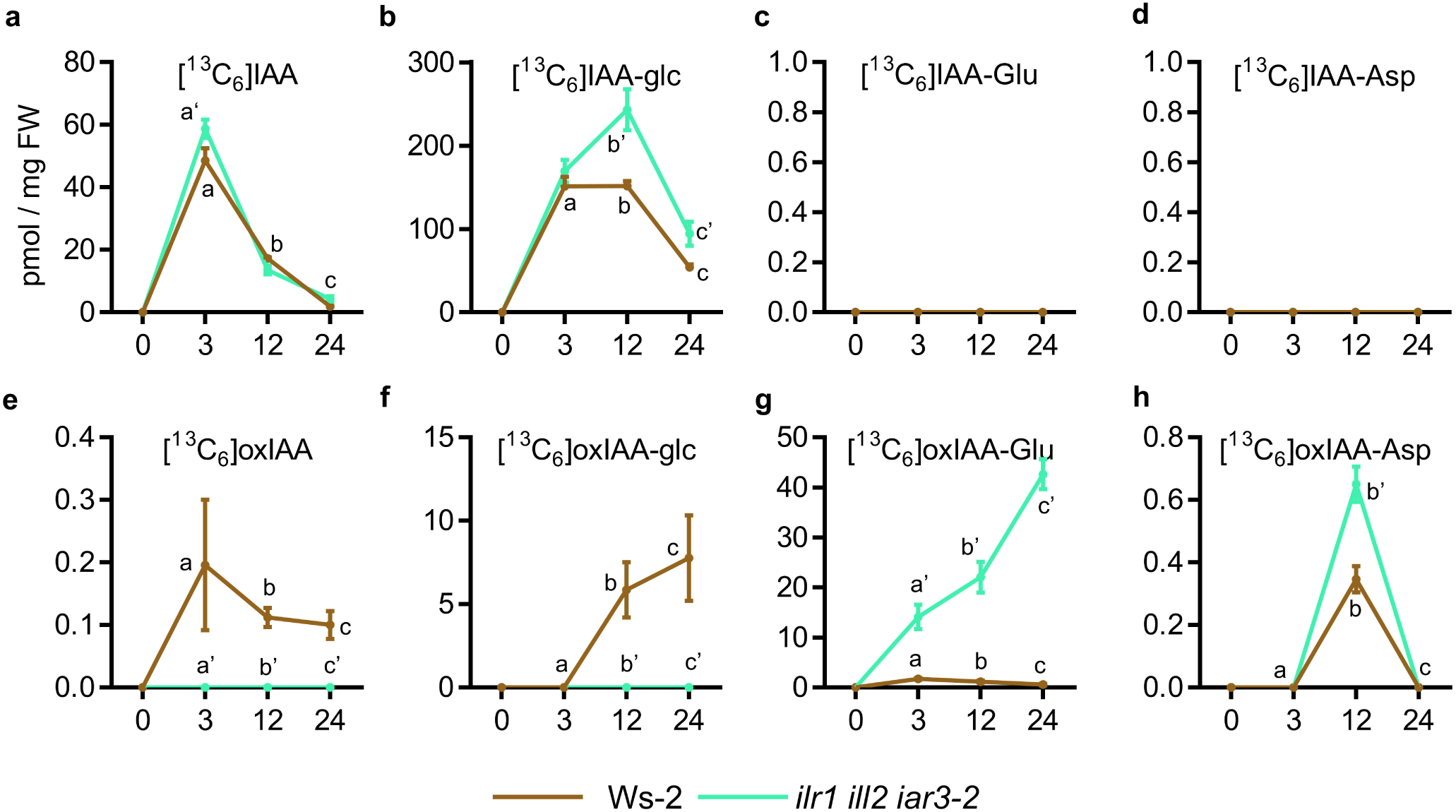
*De novo* synthesis levels of indole-3-acetic acid (IAA) metabolites in wild-type (Ws-2) and triple hydrolase *ilr1-1 iar3-2 ill2-1* mutant plants co-treated with KKI. (a-h) Formation of [^13^C_6_]-labelled IAA metabolites in 7-day-old seedlings of the indicated genotypes after incubation with 50 µM KKI in liquid media for 12 hours and then with 0.2 μM [^13^C_6_]IAA for 0, 3, 12, and 24 hours. Dots indicate the mean ± standard error of the mean. (a-h) Levels in picomoles per gram of fresh weight of (a) [^13^C_6_]IAA, (b) [^13^C_6_]IAA-glc, (c) [^13^C_6_]IAA-Glu, (d) [^13^C_6_]IAA-Asp, (e) [^13^C_6_]oxIAA, (f) [^13^C_6_]oxIAA-glc, (g) [^13^C_6_]oxIAA-Glu, and (h) [^13^C_6_]oxIAA-Asp. For each timepoint, differences were evaluated by a one-way ANOVA followed by a pairwise comparisons using a Tukeýs HSD test.

### Transcriptomics supports the auxin hyperaccumulation phenotypes in *gh3oct dao1* and implicates non-group II GH3s in the response to high IAA

We then decided to carry out a transcriptomic approach to identify the molecular pathways underlying the phenotypes and the unique responses to exogenous IAA in *gh3oct* and *gh3oct dao1-6* mutant lines. To avoid comparing organs, we used 5-day-old seedlings, as these genotypes exhibited similar plant architecture at this stage. To better understand how plants lacking a functional GH3-DAO pathway respond to increased IAA, we also included a set of samples treated with 1 µM IAA for 4 hours. The effectiveness of the IAA treatment was first confirmed by qPCR-based transcriptional induction of known IAA-responsive genes in this setup (Figure S5a), and later verified later using RNA-seq reads on the same genes (Figure S5b).

We first investigated the eight group II *GH3* genes to try to understand the potential remaining functionality in one or more of them in the *gh3oct* backgrounds. The *gh3oct* combines 8 insertions (either T-DNAs or other forms of similar transference DNA) that can generate different changes in the gene functionality. The first one is just interrupting the natural transcription of the gene. We observed this at *GH3.1* (Figure S6a), where there seems to be transcription in both sides of the insertion although there are no wild-type transcripts. Actually, by standard RNA-seq analysis (i.e. counting reads), it seems that *GH3.1* is even upregulated in the *gh3oct dao1* because the gene has not lost their IAA-responsiveness. This observation, on the other hand, furthers supports that *gh3oct dao1* accumulates more IAA than *gh3oct*.

Another usual scenario for chromatin regions with T-DNAs is the silencing of the surrounding environment. Frequently, the small RNA machinery identifies T-DNA insertions as potentially harmful sequences that have to be heterochromatinized. This is what is seen at different extents in *GH3.2*, *GH3.3, GH3.6,* and *GH3.9* (Figure S6b, c, f, g). Either before or after insertion, there appears to be a complete lack of transcription, likely driven by heterochromatinization of the region. In the tissue used for this experiment, *GH3.4* is not expressed under any condition or genotype (Figure S6d).

However, we found two insertions that are leaking some functional IAA conjugation activity because they are at the very end of the genes: *GH3.5* and *GH3.17*. While both insertions have generated a partial silencing effect, as evidenced by the signal intensity without and especially with IAA (Figure S6e, h), we cannot rule out that, at some frequency, some of these late-truncated transcripts generate at least partially functional peptides with some GH3 activity. Nevertheless, considering the different phenotypes and IAA feeding responses, we decided to continue with the transcriptomics analyses.

In absolute terms, the biggest difference in the number of differentially expressed genes (DEGs) was observed between Col-0 and *gh3oct* or *gh3oct dao1*, with more than 2,000 DEGs (Figure S7). As a reference, the IAA treatment triggered the deregulation of about 600 genes either in the wild type (Figure 7A; S7) or in the *gh3oct* and *gh3oct dao1* mutants (Figure S7). Commonly upregulated genes between *gh3oct* and *gh3oct dao1* in mock conditions were enriched in GO terms such as root meristem growth and regulation of root development of lateral root development, already pinpointing a precocious activation of lateral root developmental programs (Figure S8). Genes found to be upregulated only in *gh3oct* but not in *gh3oct dao1* were enriched in cell wall organization processes, which may also be related to lateral root organogenesis. Finally, genes found upregulated only in *gh3oct dao1* plants were highly enriched in translational-related processes (Figure S8). Downregulated genes found in both mutants were enriched for a myriad of terms related to processes such as cytoskeleton, RNA metabolism, or cell cycle (Figure S9). Particularly in *gh3oct dao1*, downregulated genes were enriched in cell division and RNA metabolism terms while no Biological Process term was enriched among the genes found downregulated only in *gh3oct* (Figure S9).

**Figure 7.**
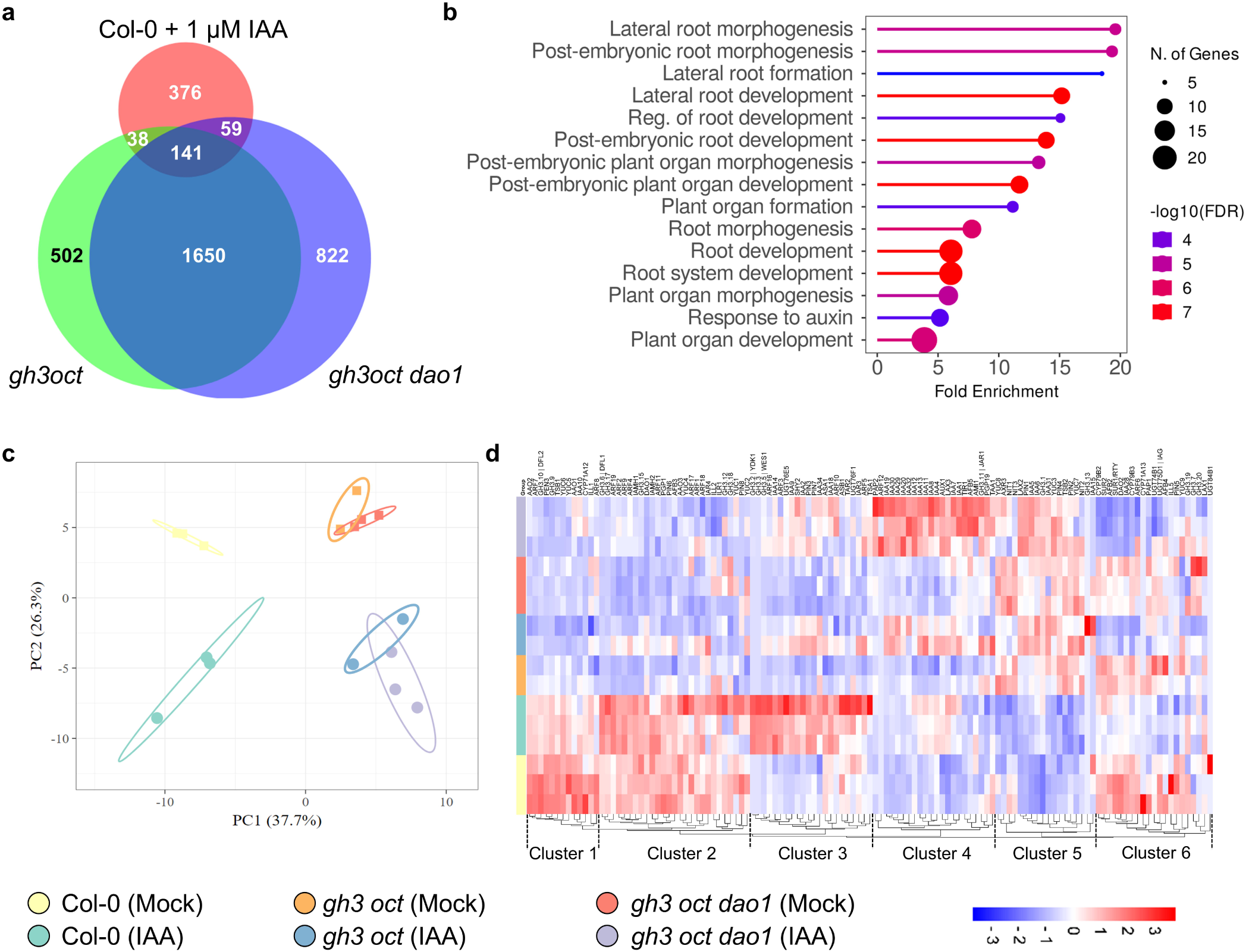
Transcriptomic analyses support IAA excess causing *gh3oct* and *gh3oct dao1* pleiotropic defects. (a) Venn diagram showing the overlapping response of IAA-treated Col-0 plants and mock-treated *gh3oct* and *gh3oct dao1* plants. (b) Lollipop chart showing the overrepresented Gene Ontology terms in the *Biological Process* category for overlapping auxin-related DEGs found in IAA-treated Col-0 plants, and mock-treated *gh3oct* and *gh3oct dao1* plants. The colour code corresponds to the –log10 of the false discovery rate (FDR), the area of the lollipop represents the number of genes in each category, and distances in the x-axis indicate the fold-enrichment of each category. (c) Principal Component Analysis of the transcriptional responses of the different genotypes and treatment samples, organized by colours. Ovals show clustering of the samples. (d) Hierarchical clustering of the transcriptional responses of the different samples. Colour scale indicates the range of normalized log2 fold-change.

Analyses of the IAA response, revealed a similar number of DEGs across all the genotypes (Figure S10, S11). GO terms were, as expected, related to auxin responses, including callus formation, wound healing, or lateral root morphogenesis (Figure S10). Genes only upregulated in one or both multiple mutants were also mainly associated with hormonal responses, including auxin and abscisic acid (Figure S10). Several genes were upregulated in Col-0 upon IAA treatment, with no response in the mutant background and were enriched in GO terms related to catabolic pathways of isoprenoids and lipids (Figure S10). Terms related to cell-cell organization were enriched among the commonly downregulated genes in response to IAA across all genotypes (Figure S11), while two specific responses were observed in *gh3oct* mutants and Col-0. In the mutants, the glucosinolate pathway appears to be downregulated, whereas in the wild type, the root hair development pathway is repressed by IAA treatment (Figure S11).

In line with our metabolic and phenotypic data, 141 DEGs were shared by IAA-treated Col-0 and mock *gh3oct* or *gh3oct dao1* plants (Figure 7a). Gene ontology analysis of these common DEGs revealed substantial enrichment for genes involved in lateral root formation, root development, or auxin response (Figure 7b). With this transcriptomic line of evidence further supporting the role of auxin accumulation underlying the mutant’s phenotypes, we took a closer look at a custom matrix of 137 auxin-related genes in the transcriptomic samples (Data S1a). A principal component analysis explained 64% of the variability among samples and identified different patterns of transcriptional misregulation, with four main clusters: (1) Col-0 mock, (2) IAA-treated Col-0, (3) both *gh3oct* and *gh3oct dao1* mock plants, and (4) IAA-treated *gh3oct* and *gh3oct dao1* genotypes (Figure 7c). We then performed hierarchical clustering of the IAA-related gene matrix for these samples, defining seven clusters (Figure 7d; Data S1b). Clusters 2 and 3 contain genes that, starting from different expression levels in the wild type, were induced upon auxin treatment, with lower induction in the mutants. Interestingly, cluster 4 grouped genes that showed mild, if any, upregulation in Col-0 upon IAA treatment, but whose induction was high in *gh3oct* and even higher *gh3oct dao1*. This enhanced induction is very likely due to the mutant plantś insufficient ability to inactivate IAA. This cluster included the auxin response factor *ARF12*; auxin transporters like *AUX1, LAX3,* and *PGP19*; auxin receptors such as *TIR1*, thirteen members of the *IAA* (*INDOLE-3-ACETIC ACID INDUCIBLE*) family, including *IAA1* (also known as *AUXIN RESISTANT 5*; *AXR5*)*, IAA4* (*AUXIN INDUCIBLE 2-11*)*, IAA7*(*AXR2*)*, IAA8, IAA11, IAA12* (*BODENLOS*)*, IAA13, IAA19* (*MASSUGU 2*)*, IAA20, IAA27* (*PAP2*)*, IAA29, IAA30,* and *IAA32* (*MEE10*) and *AFB5,* and although counterintuitive, aligned with previous modelling work^28^, the auxin biosynthesis genes *AMI1* and *TAA1* (Data S1b). Using the AGRIS tool, we further analysed the promoters of the different clusters, looking for potential enrichment in genes carrying auxin-response *cis* elements. Since this custom gene matrix is related to auxin processes, there was a general enrichment of the auxin response element. Among the clusters, clusters 3 and 4 were found to have auxin response *cis* elements in more than 77% of the genes. This observation further reinforces the idea that the genes in the cluster 4 are differentially upregulated in the mutants due to IAA excess (Data S1c).

Interestingly, cluster 4 also included *GH3.11* (*JAR1*), a non-group II GH3 member which has been shown to conjugate isoleucine to jasmonic acid (JA)^29^. This observation opened up the possibility that other non-group II GH3 members may also participate in the response to excess IAA in the absence of group II *GH3* and *DAO1*. We specifically examined the 12 non-group II *GH3* expression levels, finding that *GH3.11* and *GH3.14* (At5g13360) showed an enhanced response to IAA exclusively in the *gh3oct dao1* background. *GH3.15*, whose role in indole-butyric acid conjugation has been reported^30^, showed a higher expression in response to IAA in both the *gh3oct* and *gh3oct dao1* mutants (Figure S12). Notably, recombinant group III GH3 isoforms, mainly GH3.12, and GH3.15 and to a minor extent GH3.14, exhibited IAA-conjugating activity with Glu^21^, suggesting that the residual formation of IAA-Glu observed in the *gh3oct* mutant^19^ (Figure 5c, Figure S3b) could be ascribed to the activity of these non-group II GH3 members rather than the limited and truncated transcripts of *GH5* or *GH3.17*.

Another conspicuous trait of the *gh3oct* mutant is a strong early flowering phonotype^19^. Using a curated list of genes involved in flowering time regulation^31^ [Data S2], we analysed the effect of the *gh3oct* and *gh3oct dao1* mutations on the transcriptional output of this cohort of genes. Among the clearest candidates to explain this trait, we found a strong downregulation of the MADS-box transcription factors *FLOWERING LOCUS C*^32^, *MADS AFFECTING FLOWERING4* (*MAF4*), and *MAF5*^33^ or *TERMINAL FLOWER1*^34^, as well as derepression of *GAox1*, *GA20ox1*^35^, *AGAMOUS-LIKE14*^36^, or *REPRESSOR OF UV-B PHOTOMORPHOGENESIS 2*^37^ (Figure S13). Whether there is a direct mechanistic connection between IAA accumulation and the flowering induction remains to be determined.

## DISCUSSION

Since the identification of inactive IAA metabolites by earlier studies^12,38^, our understanding of the role of these metabolites and the pathways involved has greatly evolved, from being considered as merely static storage or waste products to the notion that IAA inactivation routes are active and regulated processes, crucial for maintaining auxin levels and spatiotemporal distribution in plants. In this study, we systematically dissected the contribution of three major inactivation routes for IAA metabolic regulation in *Arabidopsis thaliana*: oxidation via DAO enzymes, conjugation to amino acids by GH3s, and glycosylation mediated by UGTs (Figure 1), generating a comprehensive mutant toolkit combining these pathways and assessing their metabolic, developmental, and transcriptomic consequences.

Understanding of DAO-mediated IAA oxidation has undergone a significant conceptual shift since the identification of the DAO enzymes^24^. Initially, IAA was thought to be the substrate of DAOs to produce oxIAA^22-24,39^, and thus DAOs and GH3 were considered parallel and redundant pathways for IAA metabolic inactivation. Later studies showed that the product of the GH3s, IAA-amino acid conjugates, are the primary substrates for DAOs^20,25^, and further evidenced that GH3s and DAOs function in a single, linear, and major route for IAA inactivation^25^. Our [^13^C_6_]IAA feeding experiments, particularly the complete loss of *de novo* oxIAA and oxIAA-glc formation in the *ilr1 iar3 ill2* background, provide direct genetic evidence that DAO enzymes act exclusively on IAA–amino acid conjugates rather than on free IAA *in vivo*.

A major contribution of the present work concerns the functional importance of the IAA glycosylation pathway, conventionally overlooked mainly due to the absence of noticeable phenotypes in single *ugt* mutants^14-16,40^. Thus, IAA glycosylation should not be viewed as a passive storage route, but rather as a dynamically engaged buffering mechanism that limits transient auxin accumulation when the primary GH3–DAO pathway is compromised or saturated. Even though single *ugt* mutants are phenotypically silent, we show here that knocking-out both *UGT74D1* and *UGT84B1*, two functionally redundant UDP-glycosyltransferases that act on the same substrates^14,41^, results in noticeable local auxin overproduction, as evidenced by the elongated hypocotyls and the increased formation of IAA-aa conjugates in *ugt74d1 ugt84b1* seedlings. Our findings strongly indicate that IAA glycosylation significantly contributes to plant development as an active regulatory component of auxin homeostasis, while if this contribution is largely masked, it is due to functional redundancy within the UGT pathway and with other IAA inactivation pathways. The relatively mild phenotypes of *ugt74d1 ugt84b1* mutants under standard growth conditions and the more pronounced defects when in combination with *dao* mutations, however, underscore that IAA glycosylation, while impactful, plays a context-dependent role.

An intriguing aspect of our findings is the elevated IAA levels observed in the *gh3oct dao1* mutant, accompanied by both phenotypic and transcriptional signatures of auxin overaccumulation. Transcriptomic profiling indeed revealed that *gh3oct* and *gh3oct dao1* mutants considerably mimic IAA-treated wild-type plants at the gene expression level. Given that DAOs act downstream of GH3s, the enhanced auxin response in the nonuple *gh3oct dao1* mutant suggests that accumulated IAA-aa conjugates may be hydrolysed back to free IAA. This is counterintuitive, as one would expect these inactive forms to serve as a buffer or sink under high IAA conditions, rather than being mobilized. These observations imply that conjugate hydrolysis mediated by ILR1/ILL amidohydrolases primarily responds to substrate availability rather than cellular auxin status. Notably, blocking the activity of the GH3s in combination with ILR1/ILLs-mediated hydrolysis does not impair the plant’s capacity to remove excess IAA, in contrast to the observed effect when GH3s alone are knocked down. In both scenarios, however, the IAA glycosylation was upregulated. These results imply that when both the formation and breakdown of IAA-aa conjugates are blocked, the metabolic system compensates by redirecting IAA towards other inactivation pathways, likely involving alternative or yet unidentified pathways. Further research might unravel a broader metabolic flexibility in auxin inactivation than currently understood.

Although the *gh3oct* mutant has been previously used as a genetic proxy for loss of group II GH3 activity^19^, our new transcriptomics revealed that it should be considered a strong hypomorph rather than a null background. This residual activity likely reflects low-level expression of late-truncated *GH3.5* and *GH3.17* transcripts, potentially explaining previous phenotypic discrepancies between *gh3oct* and similar mutants obtained by editing^27^. Our transcriptomic analysis also identified a subset of auxin-responsive genes that was more strongly induced in the mutants than in IAA-treated wild-type plants, consistent with impaired feedback regulation due to compromised IAA inactivation capacity. Notably, this set included auxin biosynthetic genes, transporters, and numerous IAA-inducible genes, collectively indicating auxin hyperaccumulation. Interestingly, we also observed upregulation of non-group II GH3 members such as GH3.11, GH3.14 and GH3.15, suggesting a potential role for these enzymes in the response to high IAA conditions. While GH3.11 is considered highly specific for the oxylipin JA as an acyl acid substrate, thereby making a direct involvement in IAA conjugation very unlikely^42,43^, evidence exists for the group III members GH3.14, GH3.15, and GH3.12 having a relaxed substrate preference^21^. Whether these enzymes contribute to IAA conjugation *in vivo* under certain conditions will require further demonstration.

Another emerging and still enigmatic field of knowledge is the link between auxin and translation-related processes. Several lines of evidence have shown a connection between these two processes, such as the auxin-related phenotypes found in mutants for ribosomal subunits^44-47^, the translational control of some ARF2, ARF3, or ARF6 through their upstream open reading frames^48^, and the existing feedback between auxin signalling and the energetic status via the translational control exerted by TOR on the ARFs^49^. Our transcriptomic analysis of the nonuple *gh3oct dao1* mutant revealed a remarkable upregulation of translation-related GO terms, providing yet another piece of evidence for the connection auxin and translation. Additional research will be required to determine whether the accumulation of IAA alters translation generally or specifically in a subset of mRNAs.

Lastly, our transcriptomics analysis of *gh3oct* mutants also sheds light on their early-flowering phenotype. Several observations support a likely indirect role for auxin signalling in the control of flowering time. The double mutant in auxin biosynthetic genes *yuc1 yuc4,* and the polar transport mutant *pin1,* exhibits a delayed flowering transition^50^. Similarly, gain-of-function alleles of *AXR2* showed a delayed flowering time in Arabidopsis^51^, and *ARF4* knock-down lines in *Fragaria vesca* displayed a comparable delay^52^. The downregulation of floral repressors such as *FLC* or *MAF4* and *MAF5* in the IAA-accumulating *gh3oct* and *gh3oct dao1* mutants suggests that auxin overaccumulation may promote early flowering through transcriptional modulation of flowering time integrators. Whether this link is direct or indirect will require further investigation.

Collectively, our findings reveal auxin inactivation as a flexible, multi-layered metabolic network capable of redistributing flux across distinct enzymatic routes to preserve hormonal homeostasis. Our work provides a conceptual framework and genetic tools to uncover novel components of the auxin metabolic network and to understand how plants maintain hormonal balance in the face of genetic and environmental perturbations.

## MATERIALS AND METHODS

### Plant material and growth conditions

We cultured plants as in^53^. Briefly, seeds from the *Arabidopsis thaliana* (L.) Heynh. Wild types (Col-0 and Ws-2) and mutant lines were surface-sterilized with 40% v/v commercial bleach and 0.002 % Triton-X-100) for 10 minutes and then washed four times with sterile deionized water. Seeds were stratified for a minimum of 2 days and then sowed under sterile conditions on square petri dishes containing half-strength Murashige & Skoog salt mixture (M0221; Duchefa Biocemie, Haarlem, the Netherlands), 0.05% MES hydrate (M2933; Sigma), and 0.8% plant agar (P1001; Duchefa Biochemie) with the pH adjusted to 5.7 with potassium hydroxide. Plants were kept in vitro for a maximum of 2 weeks, after which they were transferred to pots containing a 3:1 mixture of organic soil and vermiculite. All plants were grown under long-day conditions (16h:8h, light:dark) at 22±1°C under cool white fluorescent light (150 µmol photons m^-2^ s^-1^). T-DNA insertional lines *dao1-1*^22^, *dao2-1*^23^, *gh3* octuple mutant^19^, *ugt74d1*^15^, and *ilr1-1 iar3-2 ill2-1*^54^, as well as the CRISPR/Cas9-generated line *ugt84b1*^14^ were previously reported.

### CRISPR/Cas9 plasmid construction

The CRISPR/Cas9-based vector to knock-out the *DAO1*-*DAO2* locus was constructed using the GreenGate system^55^ as described in^14,56^. Briefly, 4 guide RNAs (sgRNAs) targeting the *DAO1-DAO2* locus (g#7, g#4, g#6, g#3; Figure 2; Table S3) were designed using CRISPR-P (http://crispr.hzau.edu.cn/cgi-bin/CRISPR2/CRISPR). The gRNAs were generated using the primers listed in Table S1 and cloned into GreenGate D and E modules by digestion-ligation^55^. The mCherry sequence was amplified from the pGGC015 plasmid using the mCherry-*Bas*I primers (Table S1) and cloned into a B module by digestion-ligation^55^. Two supermodules were then generated by assembling the different GreenGate modules into the intermediate plasmid vectors pGGM000 and pGGN000 (Table S3). The M and N supermodules were then combined into the destination vector pGGZ003 to create the final construct (construct #1; Table S3). A separate GreenGate assembly, including 2 additional sgRNAs targeting the *DAO1* coding sequence (g#2, g#5; Figure 1; Table S3), into the destination vector pGGZ003 was performed to generate construct #2.

Additionally, a CRISPR/Cas9 construct to knock out *DAO1* (construct #3) was generated in the pKI1.1R plasmid following the protocol described in^57^. Briefly, the circular pKI1.1R plasmid was linearized by incubating 1.5 µg of the purified plasmid with the *Aar*I restriction enzyme for 10 h and then dephosphorylated using FastAP (Thermo Fisher). A target-specific gRNA was designed (g#1) using CRISPR-P 2.0 (http://crispr.hzau.edu.cn/cgi-bin/CRISPR2/CRISPR). Oligonucleotides harbouring the gRNA target (Table S1) were hybridised by slow cooling from 95-25°C and then phosphorylated using T4 polynucleotide kinase (NEB). The digested plasmid and the hybridised oligonucleotides were ligated using T4 Ligase (NEB) and then transformed into *E. coli* DH5alpha competent cells. The correct assembly of GreenGate intermediate and destination vectors was confirmed by restriction analysis. The sequence integrity in all modules and destination vectors was verified by Sanger sequencing. All constructs were mobilised into *Agrobacterium tumefaciens* GV3101 (C58C1 Rif R) cells by electroporation.

### Plant transformation and isolation of transgene-free edited mutant lines

All constructs were transferred to Arabidopsis plants by floral dipping^58^. T_1_ transgenic plants were selected on plates supplemented with 15 mg l^-1^ hygromycin B (Invitrogen).

Transformation of Col-0, *ugt74d-1*, and *gh3oct* plants with construct #1 allowed us to isolate the CRISPR/Cas9 *dao2*, *ugt74d1-1 dao1 dao2*, and *gh3oct dao2* mutant lines, respectively. The *dao1 dao2* double mutant was isolated after transformation of *dao2-1* plants with the construct #2 plasmid. Transformation of *gh3oct* plants with construct #3 led us to isolate the *gh3oct dao1* mutant. To generate the *ugt74d1 ugt84b1* mutant we transformed *ugt74d-1* plants with our previously generated CRISPR/Cas9 construct targeting *UGT84B1* (Mateo-Bonmatí *et al.*, 2021). The construct and the nature of the isolated deletions in *UGT84B1* are as previously described (Mateo-Bonmatí *et al.*, 2021). The remaining genotypes were obtained by crossing and genotyping.

### Morphometric measurements

For root and hypocotyl phenotyping, vertically grown plates were imaged using Epson Perfection V600 photo scanners. Lengths were measured from scaled images using FIJI software^59^. Differences were evaluated by a one-way ANOVA followed by pairwise comparisons using Tukeýs HSD test.

For root phenotyping in response to KKI treatment, Col-0 and *gh3oct* seeds were grown vertically in media supplemented with 0 (mock) or 5 µM KKI under long-day conditions (16 h light/8 h dark) in cultivation chambers maintained at 21 °C, with a light intensity of ∼100 μmol m−2 s−1 and 60% relative humidity. Root length and lateral root number of more than 20 seedlings per genotype were scored at 7 days after germination. Plates were imaged with a Sony α 7 II camera, and roots were quantified using ImageJ/Fiji. Differences were evaluated by a one-way ANOVA followed by pairwise comparisons using Tukeýs HSD test.

### IAA metabolite profiling

Seven-day-old *in vitro* grown seedlings (Col-0, *dao1-1*, *dao2-1*, *dao1 dao2*, *ugt84b1 ugt74d1*, *ugt74d1 dao1 dao2*, *ugt84b1 dao1 dao2*, *ugt84b1 ugt74d1 dao1 dao2*, *gh3oct*, *gh3oct dao1*, *gh3oct dao2* mutant lines) were incubated with liquid ½ MS medium containing 1 μM [^13^C_6_]IAA for 0, 3, 12, and 24 h under gentle shaking and in darkness. Seven-day-old *in vitro* grown seedlings (Ws-2, *ilr1-1 iar3-2 ill2-1*) were incubated with liquid ½ MS medium and pre-treated with 50 μM KKI for 12 h before feeding with 0.2 μM [^13^C_6_]IAA and 50 μM KKI for 0, 3, 12, and 24 h under gentle shaking and in darkness. For each time point, 10 mg whole seedlings were collected in five replicates.

Extraction, purification and quantification of targeted compounds (IAA, oxIAA, IAA-Asp, IAA-Glu, oxIAA-Asp, oxIAA-Glu, IAA-glc, and oxIAA-glc) were performed according to^60^. Briefly, samples were extracted in 1 mL of cold 50 mmol/L phosphate buffer (pH 7.0) containing 0.1% sodium diethyldithiocarbamate and a mixture of isotope-labelled internal standards. A 200 μL portion of the extract was acidified to pH 2.7 with HCl and purified using in-tip micro solid phase extraction (in-tip μSPE). Eluted samples were evaporated under reduced pressure, reconstituted in 10% aqueous methanol, and analysed using a 1290 Infinity LC system (Agilent Technologies, CA, USA) equipped with a Kinetex C18 column (50 mm x 2.1 mm, 1.7 μm; Phenomenex) and a coupled 6490 Triple Quadrupole MS system (Agilent Technologies, CA, USA).

### RT-qPCR

To test the IAA-treatment setup employed for the RNA-seq, RNA was isolated using the Total RNA Purification Kit (Norgen, Thorold, ON, Canada). DNA was removed using the RNase-Free DNase I Kit (Norgen). First-strand cDNA synthesis was performed with the iScript cDNA Synthesis Kit (Bio-Rad). The *ACTIN2* gene was used as an internal control for relative expression quantification. Four biological replicates (each being a pool of several plants) were analysed in triplicate. Quantitative PCR (qPCR) reactions were performed in 10 µl reactions containing 4 µl of LightCycler 480 SYBR Green I Master (Roche), 4 µl of PCR-grade water (Roche), 1 µl of the corresponding primer pair (10 µM each), and 1 µl of the cDNA template. The primers used are listed in Table S1. Quantification of relative gene expression was performed using the comparative *C*_T_ method (2^−ΔΔCt^)^61^ on a CFX384 Touch Real-Time PCR Detection System (Bio-Rad).

### RNA-seq

For RNA-seq, seedlings of Col-0, *gh3oct,* and *gh3oct dao1* were vertically grown on standard media in square petri dishes on top of a nylon mesh for five days. Then the nylon meshes were transferred to petri dishes with standard media (mock) or standard media supplemented with 1 µM IAA for 4 hours. Then tissue was harvested and flash-frozen. The RNA was isolated using the RNeasy Plant Mini Kit (Qiagen). RNA quality was checked by capillary electrophoresis using an Agilent 2100 Bioanalyzer. Sequencing was performed on a BGISEQ platform using PE100 reads at BGI, Hong Kong. Sequencing statistics are shown in Table S2. Clean reads were aligned to the TAIR10 reference genome with HISAT. Genotypes were analysed in triplicate, except for *gh3oct* mock and IAA treatments which could only be sequenced in duplicate because one of the libraries failed. Raw data were deposited in the Short Read Archive (SRA) under the study reference PRJNA666323.

### Bioinformatics analyses

The sgRNAs used for the CRISPR/Cas9-editing were designed using CRISPR-P (http://crispr.hzau.edu.cn/cgi-bin/CRISPR2/CRISPR). Charts shown in Figures 2-5, S1-3, S9-S10 were created using GraphPad Prism 6. All the Venn Diagrams were generated using BioVenn^62^. The PCA was created with ClustaVis^63^. The Hierarchical clustering was performed with SRplot^64^. The GO analyses and lollipop charts were built using ShinyGO^65^.

### Accession numbers

*DAO1* (AT1G14130), *DAO2* (AT1G14120), *GH3.1* (AT2G14960), *GH3.2* (AT4G37390), *GH3.3* (AT2G23170), *GH3.4* (AT1G59500), *GH3.5* (AT4G27260), *GH3.6* (AT5G54510), *GH3.9* (AT2G47750), *GH3.14* (AT5G13360), *GH3.15* (AT5G13370), *GH3.17* (AT1G28130), *ILR1* (AT3G02875), *ILL2* (AT5G56660), *IAR3* (AT1G51760) *UGT74D1* (AT2G31750), *UGT76E5* (At3g46720), *UGT84B1* (AT2G23260).

## DATA AVAILABILITY STATEMENT

All data generated or analysed during this study are provided in this published article and its supplementary data files or will be provided upon reasonable request. RNA-seq raw data were deposited in the Short Read Archive (SRA) under the study reference PRJNA666323.

## FUNDING STATEMENT

This work was partially supported by the Wallenberg Initiatives in Forest Research (WIFORCE) funded by the Knut and Alice Wallenberg Foundation (KAW 2020.0240) as well as the Swedish Governmental Agency for Innovation Systems (Vinnova); the Swedish Research Council (VR); Kempestiftelserna (JCK-1811, JCK-1111); the University of Nottingham (Nottingham Research Fellowship to U.V.) and Royal Society Research Grants (RGS_R1_191323 to U.V.); Unravelling Spatio-temporal Auxin Intracellular Redistribution for Morphogenesis (STARMORPH; 101166880 to A.P., P.H. and O.N.); TowArds Next GENeration Crops (TANGENC, no. CZ.02.01.01/00/22_008/0004581 to A.P., P.H., A.Z. and O.N.); and Grants RYC2021-030895-I (to E.M.-B.) funded by MICIU/AEI/10.13039/501100011033 and by the European Union NextGenerationEU/PRTR and PID2023-147737NA-I00 (to E.M.-B.) funded by MICIU/AEI/10.13039/501100011033 and “ERDF/EU”.

## AUTHOR CONTRUBUTIONS

R.C.-S., K.L, and E.M.-B designed the methodology. R.C.-S., and E.M.-B. obtained the multiple mutants and performed the phenotypic analyses. AŽ provided materials. R.C-S., F.B., P.H., and A.A. carried out the feeding experiments. A.P., J.Š., and O.N. performed the metabolic profiling. R.C.-S. and E.M-B. generated the RNA-seq material. E.M.-B. analysed the RNA-seq. E.M.-B, R.C.-S., U.V., M.B., O.N., and K.L analysed the data. E.M.-B., O.N., and K.L. obtained funding and provided resources. R.C.-S. and E.M.-B. prepared the original draft. All authors reviewed and edited the manuscript.

## ACKNOWLEDGEMENTS

We thank Roger Granbom (UPSC, Umeå, Sweden), and Hana Svobodová and Kateřina Perničková (LGR Olomouc, Czech Republic) for their technical assistance. We also acknowledge the Swedish Metabolomics Centre (https://www.swedishmetabolomicscentre.se/) for the access to the instrumentation.

## CONFLICT OF INTEREST

Authors declare no conflict of interest.

## SHORT LEGENDS FOR SUPPORTING INFORMATION

**Table S1.** Primer sets used in this work.

**Table S2.** Sequencing statistics of the different RNA-seq samples.

**Table S3.** Modules used for the GreenGate-based cloning of the CRISPR-Cas9 plasmids generated in this work.

**DataS1**. List of IAA-related genes, and lists of genes forming the clusters shown in Figure 6D with indication of the presence of the auxin response cis element in their corresponding promoter according to AGRIS.

**DataS2**. List of flowering time-related genes.

**Figure S1.** Phenotype of 7-day-old seedlings from newly generated mutants in different IAA inactivation pathways grown on ½ MS vertical plates.

**Figure S2.** De novo synthesis levels of indole-3-acetic acid (IAA) metabolites in assorted genotypes.

**Figure S3.** Steady-state levels of indole-3-acetic acid (IAA) metabolites in assorted genotypes.

**Figure S4.** Morphological traits of Col-0 and *gh3oct* in response to KKI.

**Figure S5.** Test of gene expression induction on the conditions we carried out the RNA-seq analysis.

**Figure S6.** *gh3oct* T-DNA insertions analysis.

**Figure S7.** Absolute number of differentially expressed genes in the different comparisons performed by RNA-seq.

**Figure S8.** Venn diagram showing the overlap between upregulated genes in gh3oct and *gh3oct dao1* in mock conditions.

**Figure S9.** Venn diagram showing the overlap between downregulated genes in *gh3oct* and *gh3oct dao1* in mock conditions.

**Figure S10.** Venn diagram showing the overlap between upregulated genes upon IAA treatment in Col-0, *gh3oct* and *gh3oct dao1*.

**Figure S11.** Venn diagram showing the overlap between downregulated genes upon IAA treatment in Col-0, *gh3oct* and *gh3oct dao1*.

**Figure S12.** Differences in the expression of non-group II *GH3* genes.

**Figure S13.** Differences in the expression of flowering time-related genes.

## Notes

### Competing Interest Statement

The authors have declared no competing interest.

## REFERENCES

1 Casanova-Sáez, R., Mateo-Bonmatí, E. & Ljung, K. Auxin Metabolism in Plants. Cold Spring Harb Perspect Biol 13, doi:10.1101/cshperspect.a039867 (2021).

2 Casanova-Sáez, R. & Voß, U. Auxin Metabolism Controls Developmental Decisions in Land Plants. Trends Plant Sci 24, 741–754, doi:10.1016/j.tplants.2019.05.006 (2019).

3 Covington, M. F. & Harmer, S. L. The circadian clock regulates auxin signaling and responses in Arabidopsis. PLoS Biol 5, e222, doi:10.1371/journal.pbio.0050222 (2007).

4 Frank, M., Cortleven, A., Pěnčík, A., Novák, O. & Schmulling, T. The Photoperiod Stress Response in Arabidopsis thaliana Depends on Auxin Acting as an Antagonist to the Protectant Cytokinin. Int J Mol Sci 23, doi:10.3390/ijms23062936 (2022).

5 Kazan, K. & Manners, J. M. Linking development to defense: auxin in plant-pathogen interactions. Trends Plant Sci 14, 373–382, doi:10.1016/j.tplants.2009.04.005 (2009).

6 Jing, H., Wilkinson, E. G., Sageman-Furnas, K. & Strader, L. C. Auxin and abiotic stress responses. J Exp Bot 74, 7000–7014, doi:10.1093/jxb/erad325 (2023).

7 Kaplinsky, N. J. & Barton, M. K. Plant biology. Plant acupuncture: sticking PINs in the right places. Science 306, 822–823, doi:10.1126/science.1105534 (2004).

8 Galweiler, L. et al. Regulation of polar auxin transport by AtPIN1 in Arabidopsis vascular tissue. Science 282, 2226–2230, doi:10.1126/science.282.5397.2226 (1998).

9 Hammes, U. Z. & Pedersen, B. P. Structure and Function of Auxin Transporters Annual Review of Plant Biology 75, 185–209 (2024).

10 Abbas, M. et al. Auxin methylation is required for differential growth in Arabidopsis. Proc Natl Acad Sci U S A 115, 6864–6869, doi:10.1073/pnas.1806565115 (2018).

11 Hall, P. J. Indole-3-acetyl-myo-inositol in kernels of *Oryza sativa*. Phytochemistry 19, 2121–2123 (1980).

12 Cohen, J. D. & Bandurski, R. S. Chemistry and Physiology of the Bound Auxins Annual Review of Plant Biology 33, 403–430 (1982).

13 Brunoni, F. et al. Conifers exhibit a characteristic inactivation of auxin to maintain tissue homeostasis. New Phytol 226, 1753–1765, doi: 10.1111/nph.16463 (2020).

14 Mateo-Bonmatí, E., Casanova-Sáez, R., Šimura, J. & Ljung, K. Broadening the roles of UDP-glycosyltransferases in auxin homeostasis and plant development. New Phytol 232, 642–654, doi:10.1111/nph.17633 (2021).

15 Tanaka, K. et al. UGT74D1 catalyzes the glucosylation of 2-oxindole-3-acetic acid in the auxin metabolic pathway in Arabidopsis. Plant Cell Physiol 55, 218–228, doi:10.1093/pcp/pct173 (2014).

16 Aoi, Y. et al. UDP-glucosyltransferase UGT84B1 regulates the levels of indole-3-acetic acid and phenylacetic acid in Arabidopsis. Biochem Biophys Res Commun 532, 244–250, doi:10.1016/j.bbrc.2020.08.026 (2020).

17 Staswick, P. E. et al. Characterization of an Arabidopsis enzyme family that conjugates amino acids to indole-3-acetic acid. Plant Cell 17, 616–627, doi:10.1105/tpc.104.026690 (2005).

18 Staswick, P. E., Tiryaki, I. & Rowe, M. L. Jasmonate response locus JAR1 and several related Arabidopsis genes encode enzymes of the firefly luciferase superfamily that show activity on jasmonic, salicylic, and indole-3-acetic acids in an assay for adenylation. Plant Cell 14, 1405–1415, doi:10.1105/tpc.000885 (2002).

19 Casanova-Sáez, R. et al. Inactivation of the entire Arabidopsis group II GH3s confers tolerance to salinity and water deficit. New Phytol 235, 263–275, doi:10.1111/nph.18114 (2022).

20 Müller, K. et al. DIOXYGENASE FOR AUXIN OXIDATION 1 catalyzes the oxidation of IAA amino acid conjugates. Plant Physiol 187, 103–115, doi:10.1093/plphys/kiab242 (2021).

21 Brunoni, F. et al. Amino acid conjugation of oxIAA is a secondary metabolic regulation involved in auxin homeostasis. New Phytol 238, 2264–2270, doi:10.1111/nph.18887 (2023).

22 Porco, S. et al. Dioxygenase-encoding AtDAO1 gene controls IAA oxidation and homeostasis in Arabidopsis. Proc Natl Acad Sci U S A 113, 11016–11021, doi:10.1073/pnas.1604375113 (2016).

23 Zhang, J. et al. DAO1 catalyzes temporal and tissue-specific oxidative inactivation of auxin in Arabidopsis thaliana. Proc Natl Acad Sci U S A 113, 11010–11015, doi:10.1073/pnas.1604769113 (2016).

24 Zhao, Z. et al. A role for a dioxygenase in auxin metabolism and reproductive development in rice. Dev Cell 27, 113–122, doi:10.1016/j.devcel.2013.09.005 (2013).

25 Hayashi, K. I. et al. The main oxidative inactivation pathway of the plant hormone auxin. Nat Commun 12, 6752, doi:10.1038/s41467-021-27020-1 (2021).

26 Fukui, K. et al. Chemical inhibition of the auxin inactivation pathway uncovers the roles of metabolic turnover in auxin homeostasis. Proc Natl Acad Sci U S A 119, e2206869119, doi:10.1073/pnas.2206869119 (2022).

27 Guo, R. et al. Local conjugation of auxin by the GH3 amido synthetases is required for normal development of roots and flowers in Arabidopsis. Biochem Biophys Res Commun 589, 16–22, doi:10.1016/j.bbrc.2021.11.109 (2022).

28 Mellor, N. et al. Dynamic regulation of auxin oxidase and conjugating enzymes AtDAO1 and GH3 modulates auxin homeostasis. Proc Natl Acad Sci U S A 113, 11022–11027, doi:10.1073/pnas.1604458113 (2016).

29 Suza, W. P. & Staswick, P. E. The role of JAR1 in Jasmonoyl-L: -isoleucine production during Arabidopsis wound response. Planta 227, 1221–1232, doi:10.1007/s00425-008-0694-4 (2008).

30 Sherp, A. M., Westfall, C. S., Alvarez, S. & Jez, J. M. *Arabidopsis thaliana* GH3.15 acyl acid amido synthetase has a highly specific substrate preference for the auxin precursor indole-3-butyric acid. J Biol Chem 293, 4277–4288, doi:10.1074/jbc.RA118.002006 (2018).

31 Bouche, F., Lobet, G., Tocquin, P. & Perilleux, C. FLOR-ID: an interactive database of flowering-time gene networks in *Arabidopsis thaliana*. Nucleic Acids Res 44, 1167–1171, doi:10.1093/nar/gkv1054 (2016).

32 Michaels, S. D. & Amasino, R. M. *FLOWERING LOCUS C* encodes a novel MADS domain protein that acts as a repressor of flowering. Plant Cell 11, 949–956, doi:10.1105/tpc.11.5.949 (1999).

33 Kim, D. H. & Sung, S. Coordination of the vernalization response through a VIN3 and FLC gene family regulatory network in Arabidopsis. Plant Cell 25, 454–469, doi:10.1105/tpc.112.104760 (2013).

34 Shannon, S. & Meeks-Wagner, D. R. A Mutation in the Arabidopsis TFL1 Gene Affects Inflorescence Meristem Development. Plant Cell 3, 877–892, doi:10.1105/tpc.3.9.877 (1991).

35 Coles, J. P. et al. Modification of gibberellin production and plant development in Arabidopsis by sense and antisense expression of gibberellin 20-oxidase genes. Plant J 17, 547–556, doi:10.1046/j.1365-313x.1999.00410.x (1999).

36 Perez-Ruiz, R. V. et al. XAANTAL2 (AGL14) Is an Important Component of the Complex Gene Regulatory Network that Underlies Arabidopsis Shoot Apical Meristem Transitions. Mol Plant 8, 796–813, doi:10.1016/j.molp.2015.01.017 (2015).

37 Wang, W., Yang, D. & Feldmann, K. A. EFO1 and EFO2, encoding putative WD-domain proteins, have overlapping and distinct roles in the regulation of vegetative development and flowering of Arabidopsis. J Exp Bot 62, 1077–1088, doi:10.1093/jxb/erq336 (2011).

38 Östin, A., Kowalyczk, M., Bhalerao, R. P. & Sandberg, G. Metabolism of indole-3-acetic acid in Arabidopsis. Plant Physiol 118, 285–296, doi:10.1104/pp.118.1.285 (1998).

39 Takehara, S. et al. A common allosteric mechanism regulates homeostatic inactivation of auxin and gibberellin. Nat Commun 11, 2143, doi:10.1038/s41467-020-16068-0 (2020).

40 Škyvarová, D., Brunoni, F., Žukauskaitė, A. & Pěnčík, A. Glycosylation pathways in auxin homeostasis. Physiologia Plantarum 177, e70170 (2025).

41 Brunoni, F. et al. A bacterial assay for rapid screening of IAA catabolic enzymes. Plant Methods 15, 126, doi:10.1186/s13007-019-0509-6 (2019).

42 Holland, C. K. & Jez, J. M. Fidelity in plant hormone modifications catalyzed by Arabidopsis GH3 acyl acid amido synthetases. Journal of Biological Chemistry 300, 107421 (2024).

43 Široká, J. et al. Amide conjugates of the jasmonate precursor cis-(+)-12-oxo-phytodienoic acid regulate its homeostasis during plant stress responses. Plant Physiology 197, kiae636 (2025).

44 Nishimura, T., Wada, T. & Okada, K. A key factor of translation reinitiation, ribosomal protein L24, is involved in gynoecium development in Arabidopsis. Biochem Soc Trans 32, 611–613, doi:10.1042/BST0320611 (2004).

45 Zhou, F., Roy, B. & von Arnim, A. G. Translation reinitiation and development are compromised in similar ways by mutations in translation initiation factor eIF3h and the ribosomal protein RPL24. BMC Plant Biol 10, 193, doi:10.1186/1471-2229-10-193 (2010).

46 Rosado, A. et al. Auxin-mediated ribosomal biogenesis regulates vacuolar trafficking in Arabidopsis. Plant Cell 22, 143–158, doi:10.1105/tpc.109.068320 (2010).

47 Rosado, A. & Raikhel, N. V. Application of the gene dosage balance hypothesis to auxin-related ribosomal mutants in Arabidopsis. Plant Signal Behav 5, 450–452, doi:10.4161/psb.5.4.11341 (2010).

48 Rosado, A., Li, R., van de Ven, W., Hsu, E. & Raikhel, N. V. Arabidopsis ribosomal proteins control developmental programs through translational regulation of auxin response factors. Proc Natl Acad Sci U S A 109, 19537–19544, doi:10.1073/pnas.1214774109 (2012).

49 Schepetilnikov, M. et al. TOR and S6K1 promote translation reinitiation of uORF-containing mRNAs via phosphorylation of eIF3h. EMBO J 32, 1087–1102, doi:10.1038/emboj.2013.61 (2013).

50 Cheng, Y., Dai, X. & Zhao, Y. Auxin synthesized by the YUCCA flavin monooxygenases is essential for embryogenesis and leaf formation in Arabidopsis. Plant Cell 19, 2430–2439, doi:10.1105/tpc.107.053009 (2007).

51 Mai, Y. X., Wang, L. & Yang, H. Q. A gain-of-function mutation in IAA7/AXR2 confers late flowering under short-day light in Arabidopsis. J Integr Plant Biol 53, 480–492, doi:10.1111/j.1744-7909.2011.01050.x (2011).

52 Dong, X. et al. Auxin-induced AUXIN RESPONSE FACTOR4 activates APETALA1 and FRUITFULL to promote flowering in woodland strawberry. Hortic Res 8, 115, doi:10.1038/s41438-021-00550-x (2021).

53 Casanova-Sáez, R. et al. A suitable strategy to find IAA metabolism mutants. Physiol Plant 177, e70166, doi:10.1111/ppl.70166 (2025).

54 Rampey, R. A. et al. A Family of Auxin-Conjugate Hydrolases That Contributes to Free Indole-3-Acetic Acid Levels during Arabidopsis Germination. Plant Physiology 135, 978–988 (2004).

55 Lampropoulos, A. et al. GreenGate---a novel, versatile, and efficient cloning system for plant transgenesis. PLoS One 8, e83043, doi:10.1371/journal.pone.0083043 (2013).

56 Capovilla, G., Symeonidi, E., Wu, R. & Schmid, M. Contribution of major FLM isoforms to temperature-dependent flowering in Arabidopsis thaliana. J Exp Bot 68, 5117–5127, doi:10.1093/jxb/erx328 (2017).

57 Tsutsui, H. & Higashiyama, T. pKAMA-ITACHI Vectors for Highly Efficient CRISPR/Cas9-Mediated Gene Knockout in Arabidopsis thaliana. Plant Cell Physiol 58, 46–56, doi:10.1093/pcp/pcw191 (2017).

58 Clough, S. J. & Bent, A. F. Floral dip: a simplified method for Agrobacterium-mediated transformation of Arabidopsis thaliana. Plant J 16, 735–743, doi:10.1046/j.1365-313x.1998.00343.x (1998).

59 Schindelin, J. et al. Fiji: an open-source platform for biological-image analysis. Nat Methods 9, 676–682, doi:10.1038/nmeth.2019 (2012).

60 Hladík, P., Petřík, I., Žukauskaité, A., Novák, O. & Pěnčík, A. Metabolic profiles of 2-oxindole-3-acetyl-amino acid conjugates differ in various plant species. Front Plant Sci 14, 1217421, doi:10.3389/fpls.2023.1217421 (2023).

61 Schmittgen, T. D. & Livak, K. J. Analyzing real-time PCR data by the comparative C(T) method. Nat Protoc 3, 1101–1108, doi:10.1038/nprot.2008.73 (2008).

62 Hulsen, T., de Vlieg, J. & Alkema, W. BioVenn - a web application for the comparison and visualization of biological lists using area-proportional Venn diagrams. BMC Genomics 9, 488, doi:10.1186/1471-2164-9-488 (2008).

63 Metsalu, T. & Vilo, J. ClustVis: a web tool for visualizing clustering of multivariate data using Principal Component Analysis and heatmap. Nucleic Acids Res 43, 566–570, doi:10.1093/nar/gkv468 (2015).

64 Tang, D. et al. SRplot: A free online platform for data visualization and graphing. PLoS One 18, e0294236, doi:10.1371/journal.pone.0294236 (2023).

65 Ge, S. X., Jung, D. & Yao, R. ShinyGO: a graphical gene-set enrichment tool for animals and plants. Bioinformatics 36, 2628–2629, doi:10.1093/bioinformatics/btz931 (2020).

