## Supplementary Figures for "Comprehensive characterisation of IAA inactivation pathways reveals the impact of glycosylation on auxin metabolism and plant development"

Supplementary Information

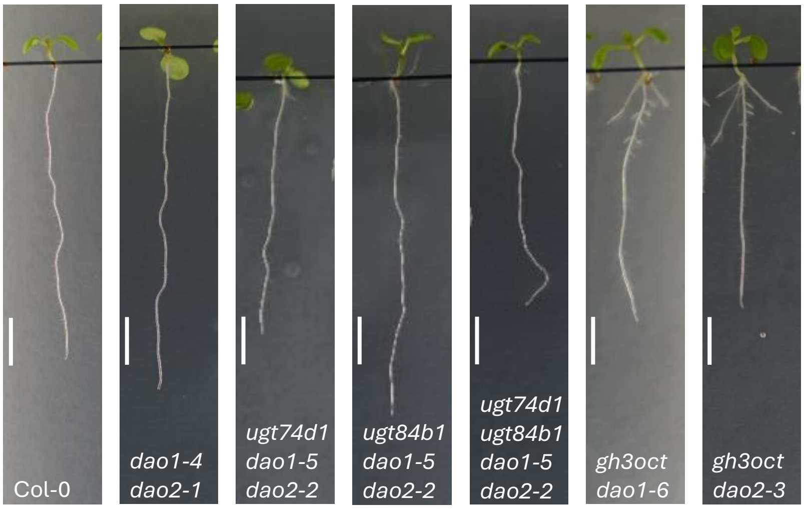


**Figure S1**. Phenotype of 7-day-old seedlings from newly generated mutants in different IAA inactivation pathways grown on ½ MS vertical plates. Scale bars represent 0.5 cm.

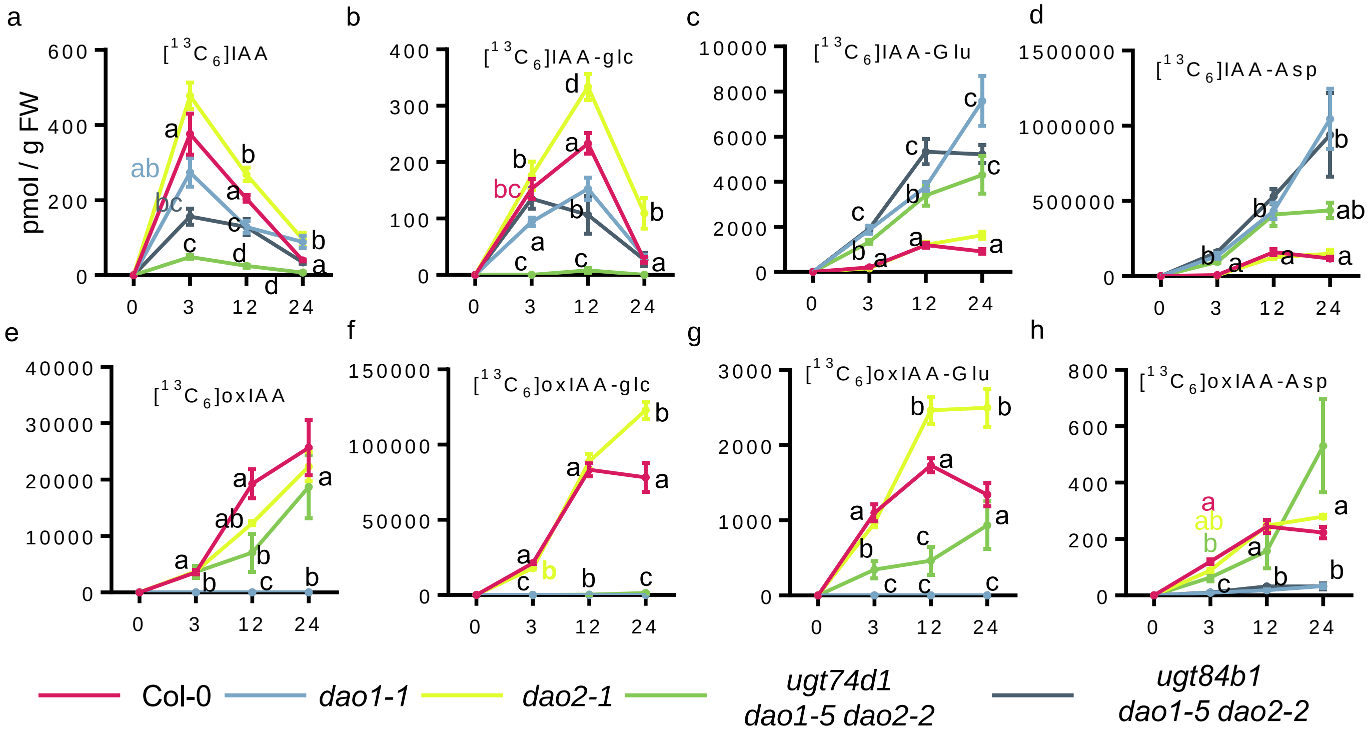


**Figure S2**. Levels of *de novo* synthesized indole-3-acetic acid (IAA) metabolites in assorted genotypes.
(A-H) Formation of [$^{13}C_6$]-labelled IAA metabolites in 7-day-old seedlings of the indicated genotypes after incubation with [$^{13}C_6$]IAA for 0, 3, 12, and 24 hours. Dots indicate the mean ± standard error of the mean. Levels in picomoles per gram of fresh weight of (a) [$^{13}C_6$]IAA, (b) [$^{13}C_6$]IAA-glc, (c) [$^{13}C_6$]IAA-Glu, (d) [$^{13}C_6$]IAA-Asp, (e) [$^{13}C_6$]oxIAA, (f) [$^{13}C_6$]oxIAA-glc, (g) [$^{13}C_6$]oxIAA-Glu, and (h) [$^{13}C_6$]oxIAA-Asp. For each time point, differences were evaluated by one-way ANOVA followed by pairwise comparisons using Tukey´s HSD test.

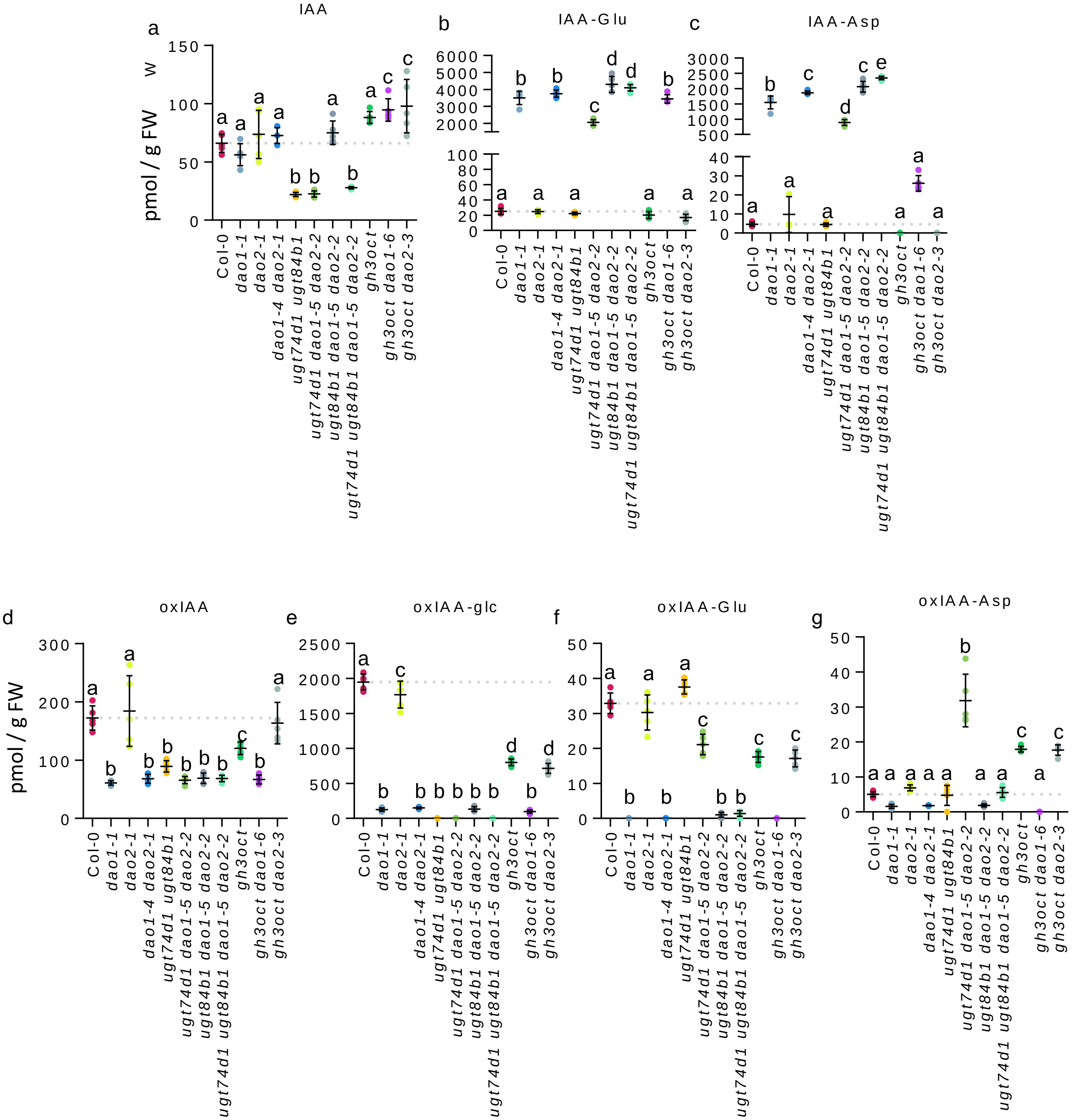


**Figure S3**. Steady-state levels of indole-3-acetic acid (IAA) metabolites in assorted genotypes. Plant samples consisted of roots from 5-day-old seedlings. Dots indicate the individual values of replicates. Error bars represent the standard error of the mean. Levels of (a) IAA, (b) IAA-Glu, (c) IAA-Asp, (d) oxIAA, (e) oxIAA-glc, (f) oxIAA-Glu, and (g) oxIAA-Asp are expressed in picomoles per gram of fresh weight. Levels of IAA-glc could not be reliably measured in these samples due to an unknown co-eluting component masking the detection of the IAA-glc peak. Differences were evaluated by one-way ANOVA followed by pairwise comparisons using Tukey´s HSD test.

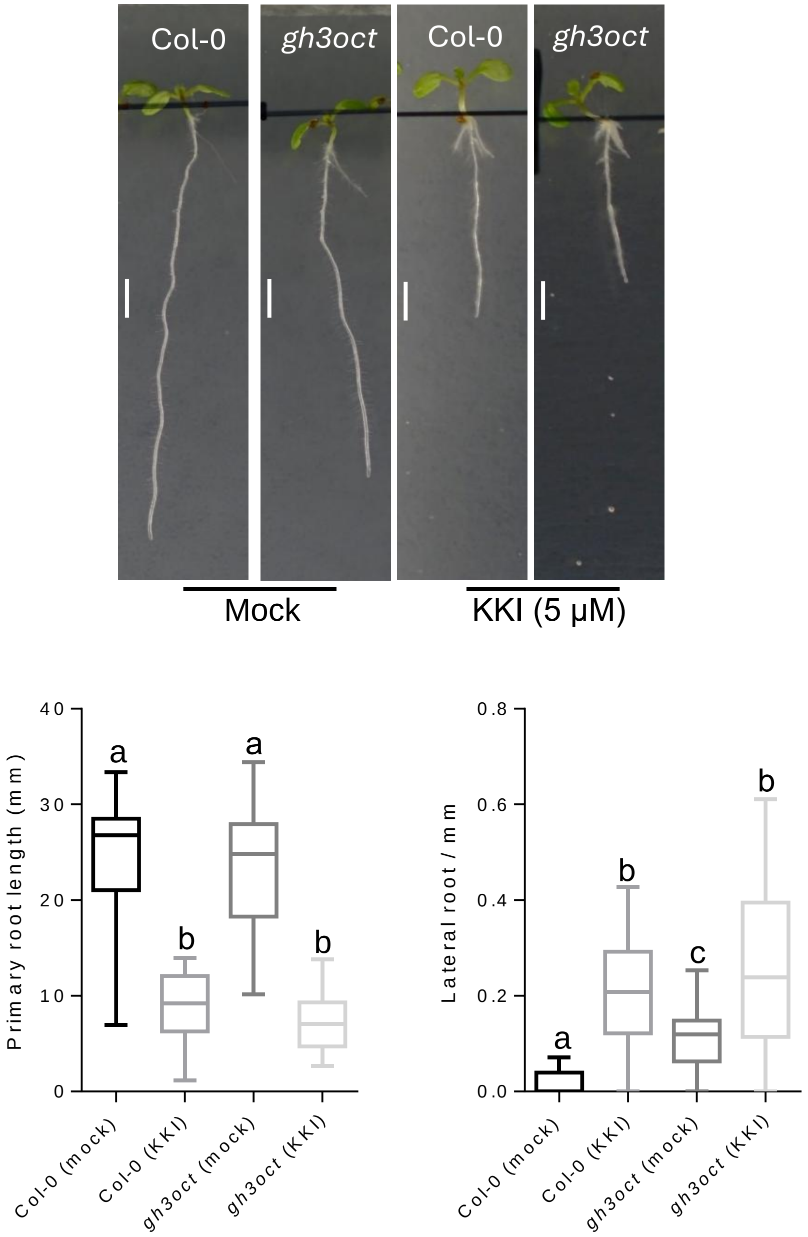


**Figure S4. Morphological traits of Col-0 and *gh3oct* in response to KKI.** Representative pictures of Col-0 and *gh3oct* grown in mock and 5 µM of KKI. Primary root length and lateral root density measurements of the above-mentioned genotypes and treatments. Differences were evaluated by one-way ANOVA followed by pairwise comparisons using Tukey´s HSD test (n>23).

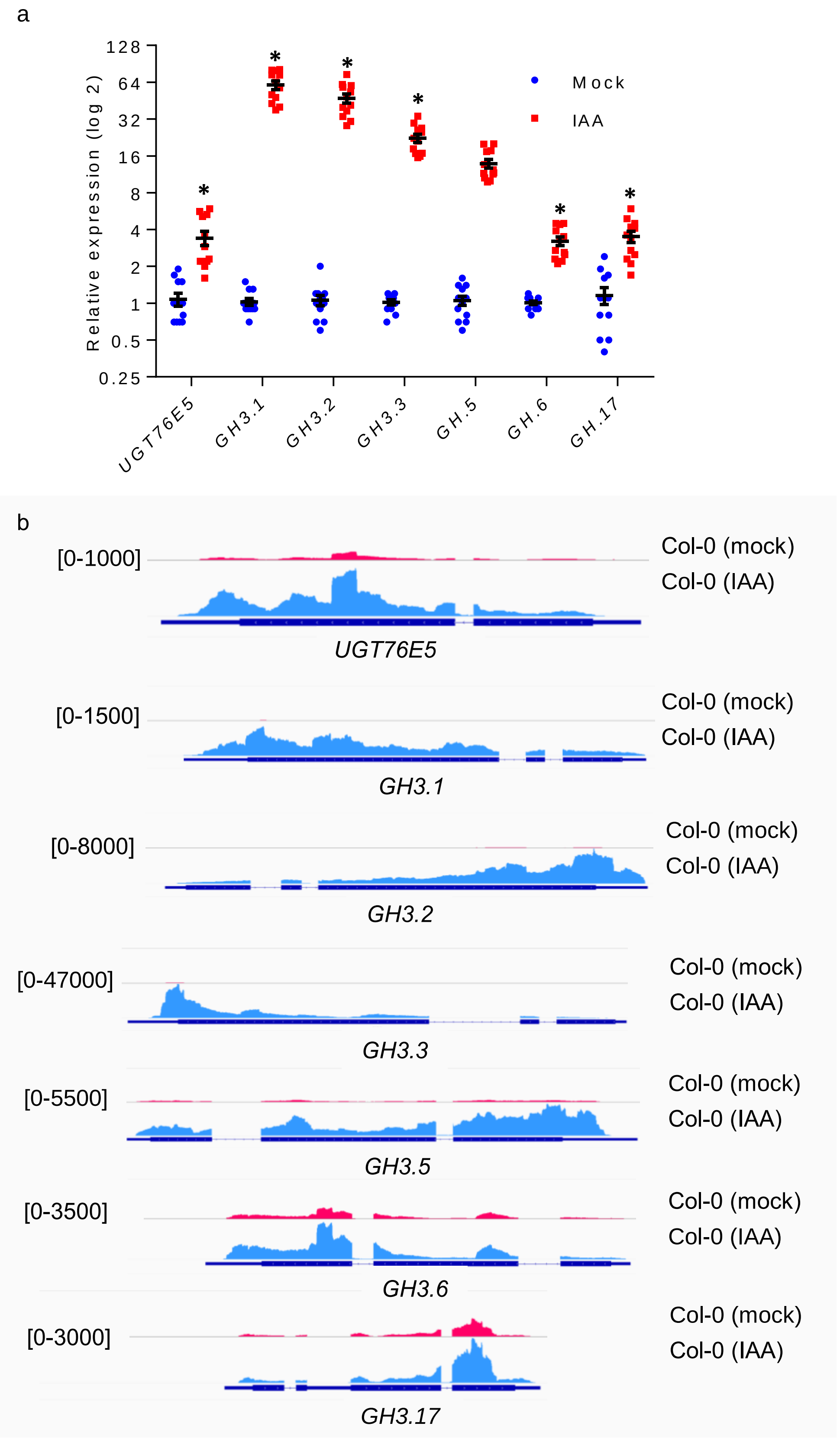


**Figure S5**. Test of gene expression induction under the conditions we carried out the RNA-seq analysis. (a) Relative expression values of the IAA-responsive genes *UGT76E5*, *GH3.1*, *GH3.2*, *GH3.3*, *GH3.5*, *GH3.6*, and *GH3.17* using Col-0 plants grown for five days on a nylon mesh and transferred to another plate with 1 μM IAA for 4 hours. Values were normalized to the mock value. Asterisk indicates statistically significant differences to mock treatment (* $p < 0.0001$; n = 12). (b) Screenshots of RNA-seq profiles over the same genes shown in (a) in the transcriptomics experiment. Profiles were visualized on IGV and scaled to the signal on IAA-treated samples. The upper and bottom panels correspond to Col-0 mock and IAA-treated, respectively.

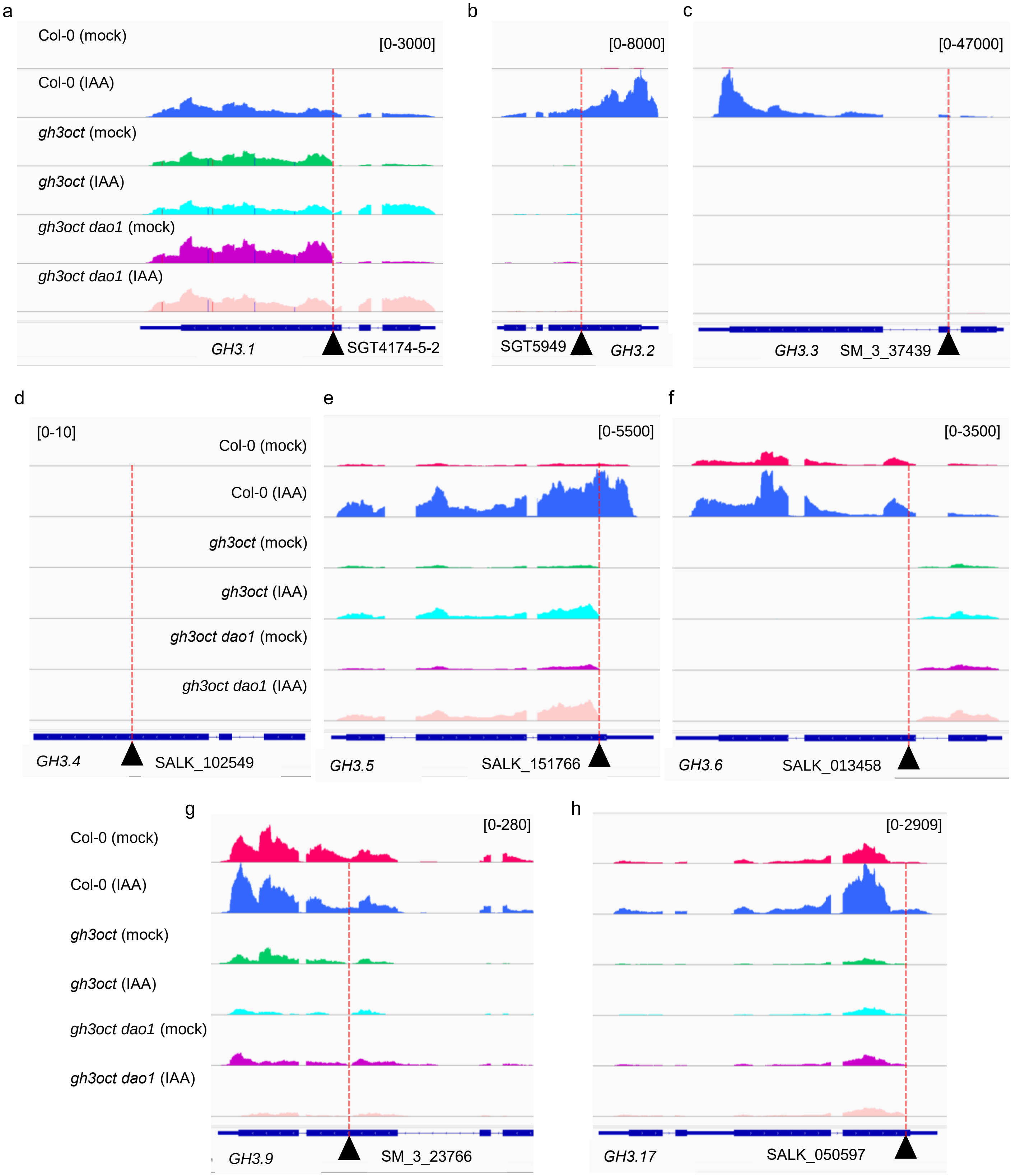


**Figure S6**. *gh3oct* T-DNA insertions analysis. (a-h) IGV tracks of Col-0 (mock), Col-0 (treated with IAA), *gh3oct* (mock), *gh3oct* (treated with IAA), *gh3oct dao1-6* (mock), and *gh3oct dao1-6* (treated with IAA), represented in different colours of (a) *GH3.1*, (b) *GH3.2*, (c) *GH3.3*, (d) *GH3.4*, (e) *GH3.5*, (f) *GH3.6*, (g) *GH3.9*, and (h) *GH3.17*. The bottom tracks corresponds to the gene annotation. Triangles represent the insertion present in each gene. A red dashed line marks the insertion point. Colour vertical lines present in panel (a) correspond to reads carrying polymorphisms between Col-0 (reference genome) and L*er* (original background of some of the insertional alleles).

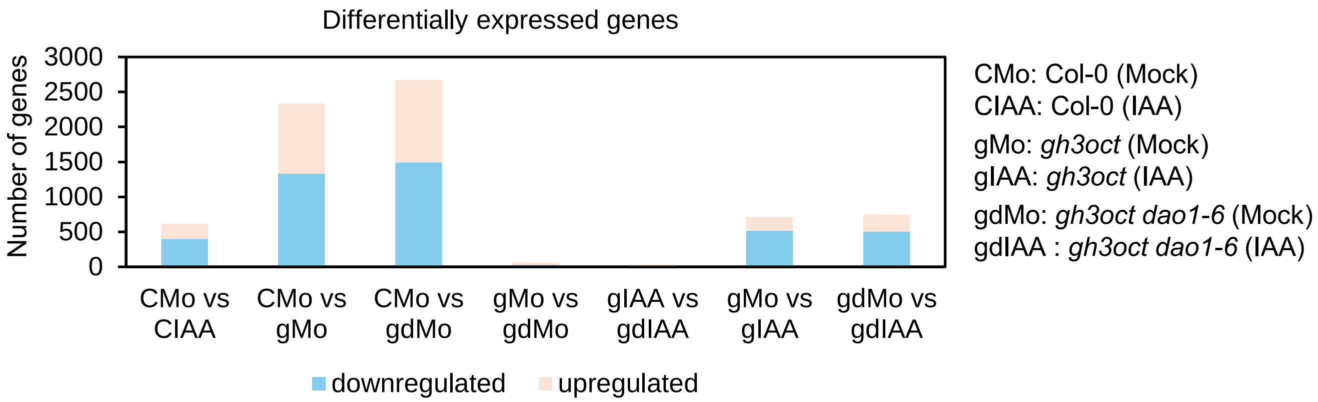


**Figure S7**. Absolute number of differentially expressed genes in the different comparisons performed by RNA-seq.

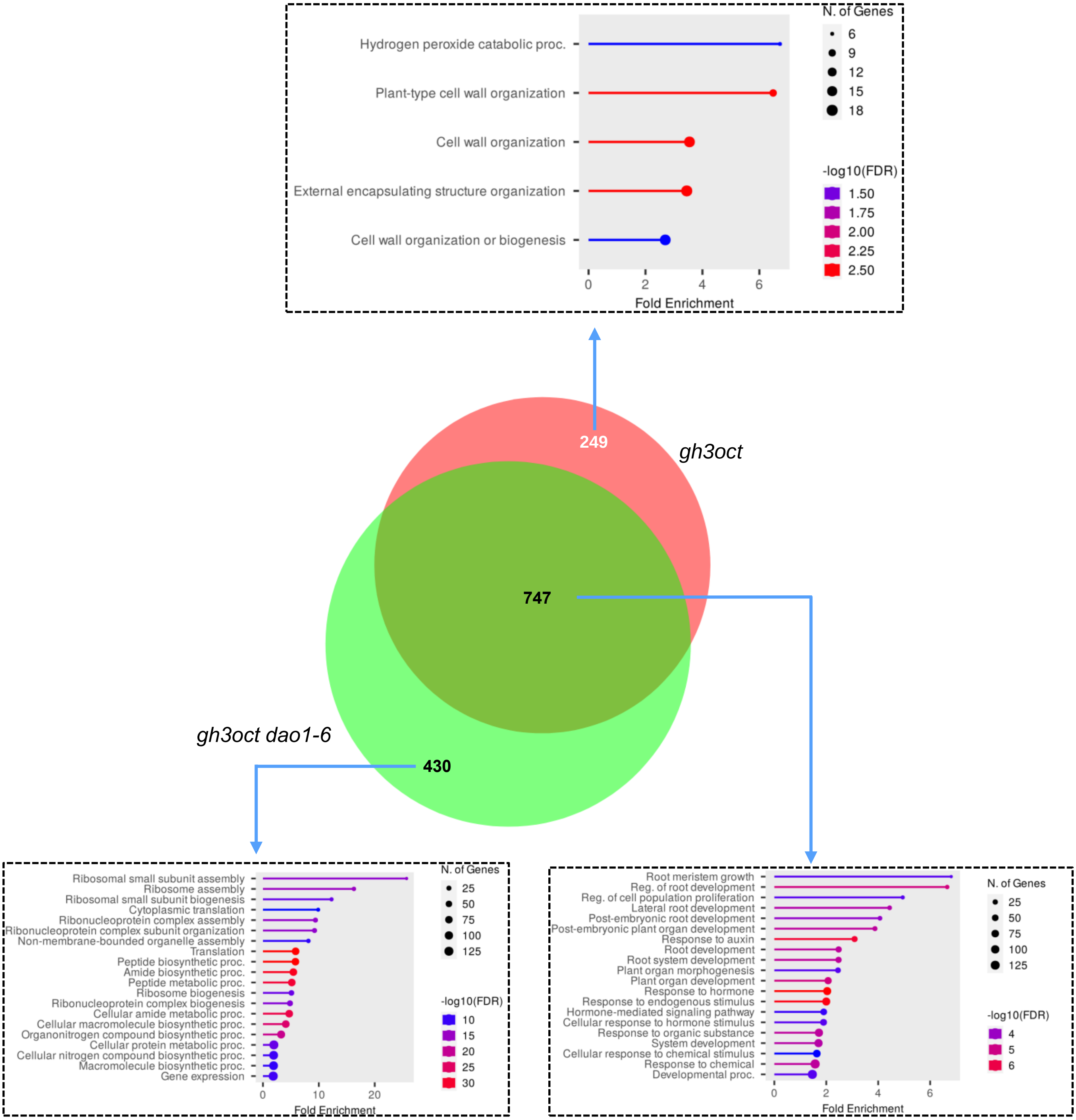


**Figure S8**. Venn diagram showing the overlap between upregulated genes in *gh3oct* and *gh3oct dao1* in mock conditions. Overrepresented Gene Ontology (GO) terms corresponding to the *Biological Process* category are shown in lollipop charts. Lollipop charts were created with Shiny-GO using default parameters. The colour code corresponds to the –log10 of the false discovery rate (FDR), the area of the lollipop represents the number of genes in each category, and distances in the x-axis indicate the fold-enrichment of each category.

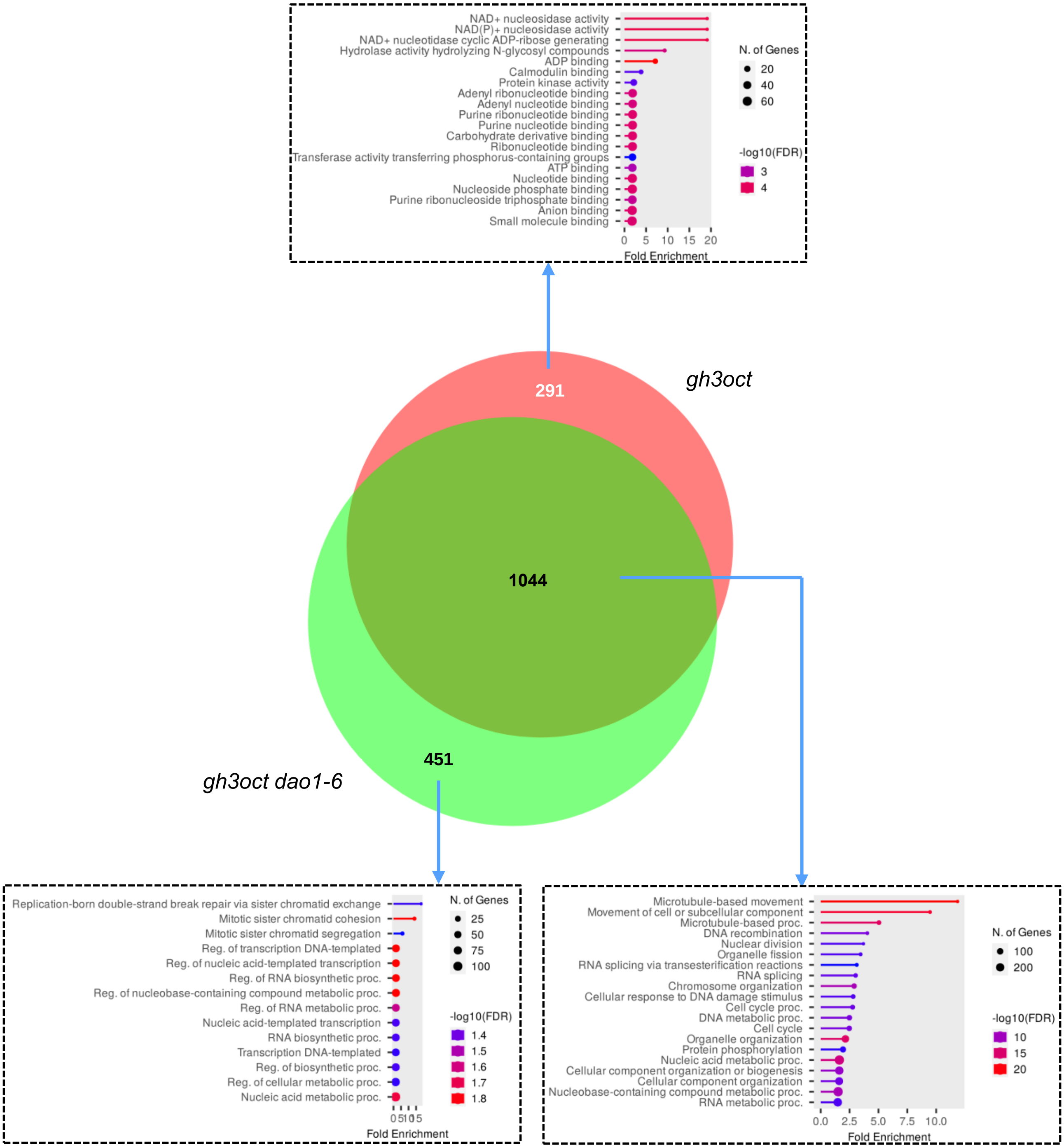


**Figure S9**. Venn diagram showing the overlap between downregulated genes in *gh3oct* and *gh3oct dao1* in mock conditions. Overrepresented Gene Ontology (GO) terms corresponding to the *Biological Process* (bottom panels) and *Molecular function* (top panel*)* categories are shown in lollipop charts. Lollipop charts were created with Shiny-GO using default parameters. The colour code corresponds to the –log10 of the false discovery rate (FDR), the area of the lollipop represents the number of genes in each category, and distances in the x-axis indicate the fold-enrichment of each category.

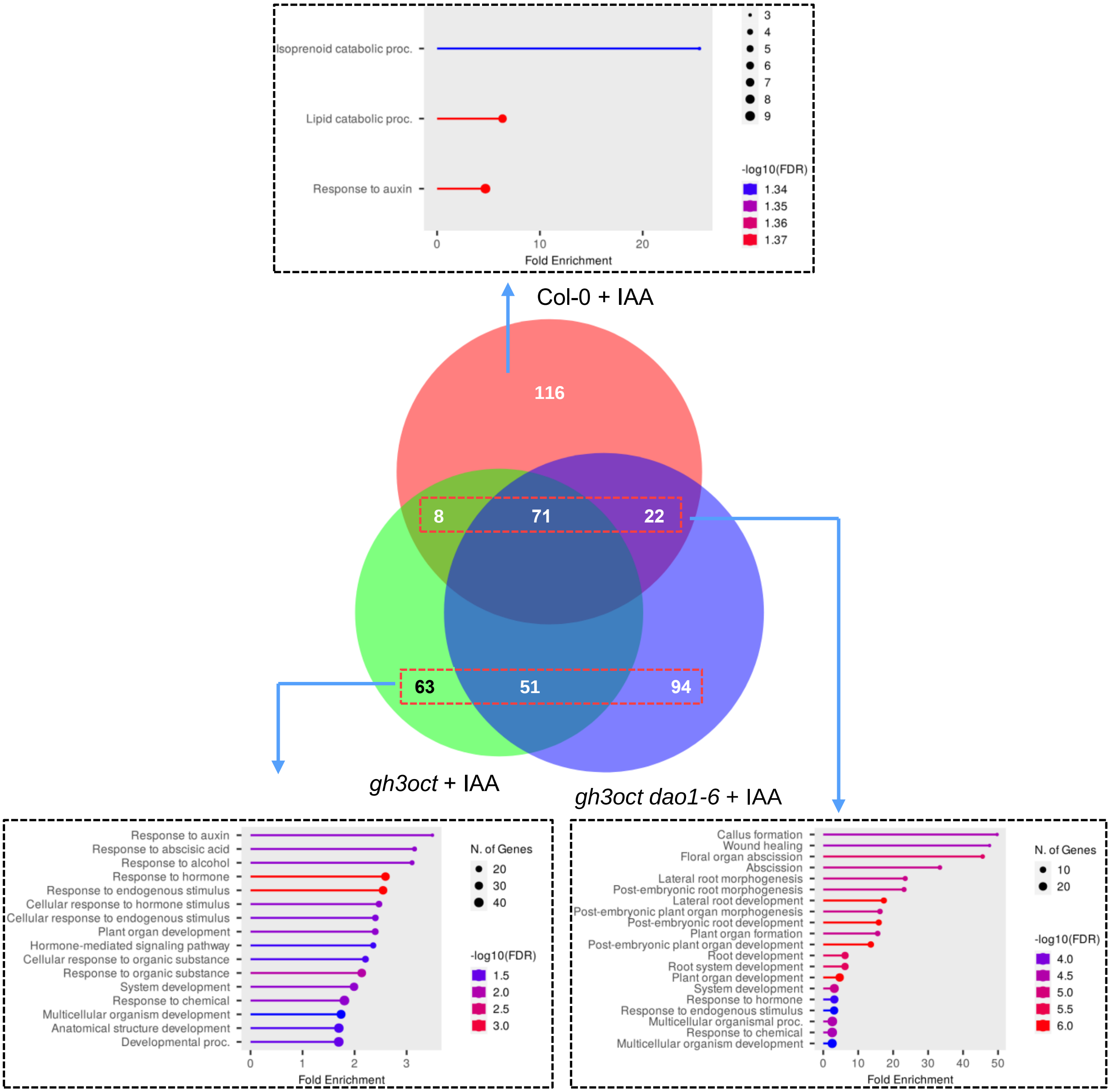


**Figure S10.** Venn diagram showing the overlap between upregulated genes upon IAA treatment in Col-0, *gh3oct* and *gh3oct dao1*. Overrepresented Gene Ontology (GO) terms corresponding to the *Biological Process* category are shown in lollipop charts. Lollipop charts were created with Shiny-GO using default parameters. The colour code corresponds to the –log10 of the false discovery rate (FDR), the area of the lollipop represents the number of genes in each category, and distances in the x-axis indicate the fold-enrichment of each category.

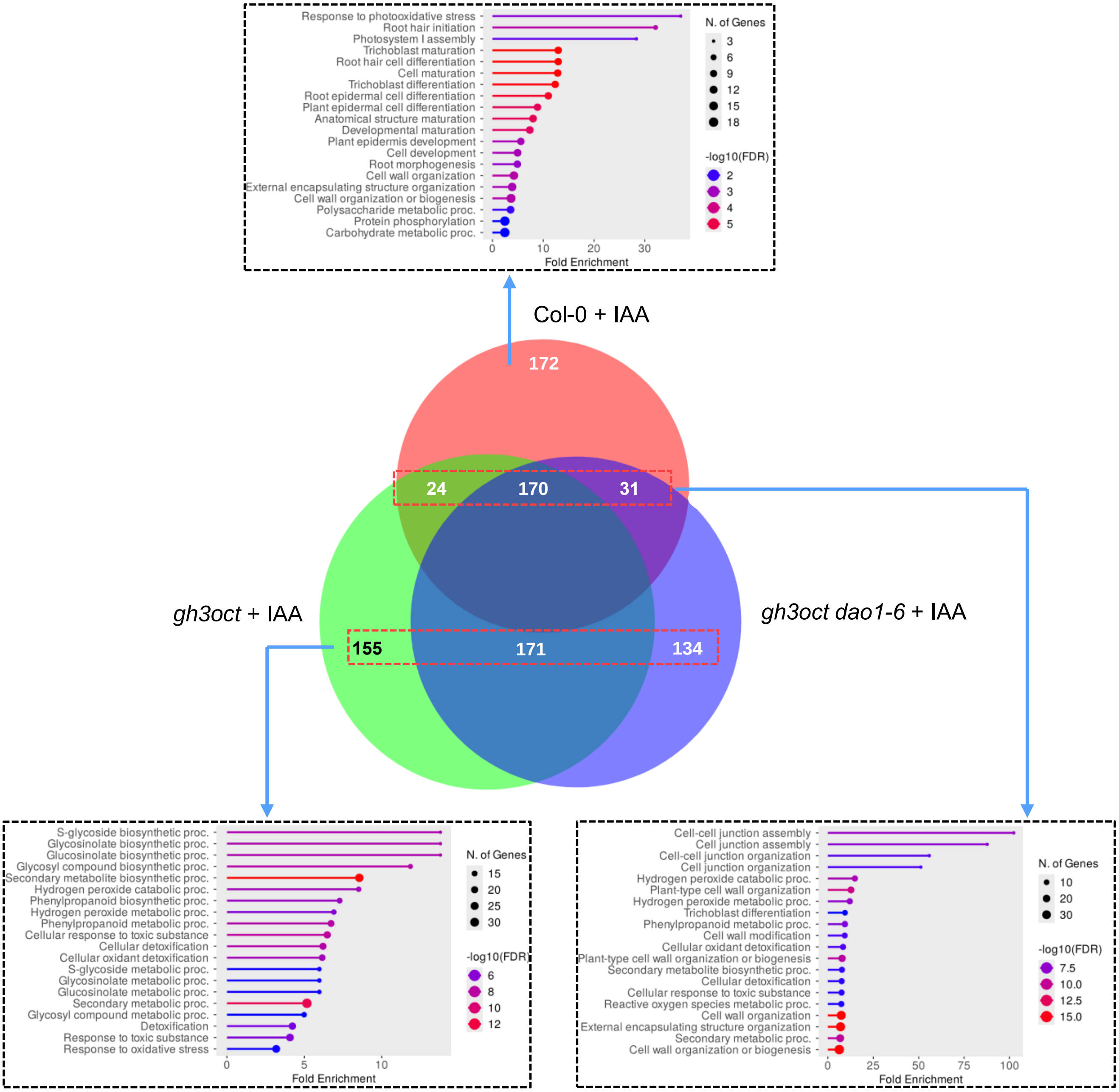


**Figure S11**. Venn diagram showing the overlap between downregulated genes upon IAA treatment in Col-0, *gh3oct* and *gh3oct dao1*. Overrepresented Gene Ontology (GO) terms corresponding to the *Biological Process* category are shown in lollipop charts. Lollipop charts were created with Shiny-GO using default parameters. The colour code corresponds to the –log10 of the false discovery rate (FDR), the area of the lollipop represents the number of genes in each category, and distances in the x-axis indicate the fold-enrichment of each category.

Casanova-Sáez *et al.* Figure S12

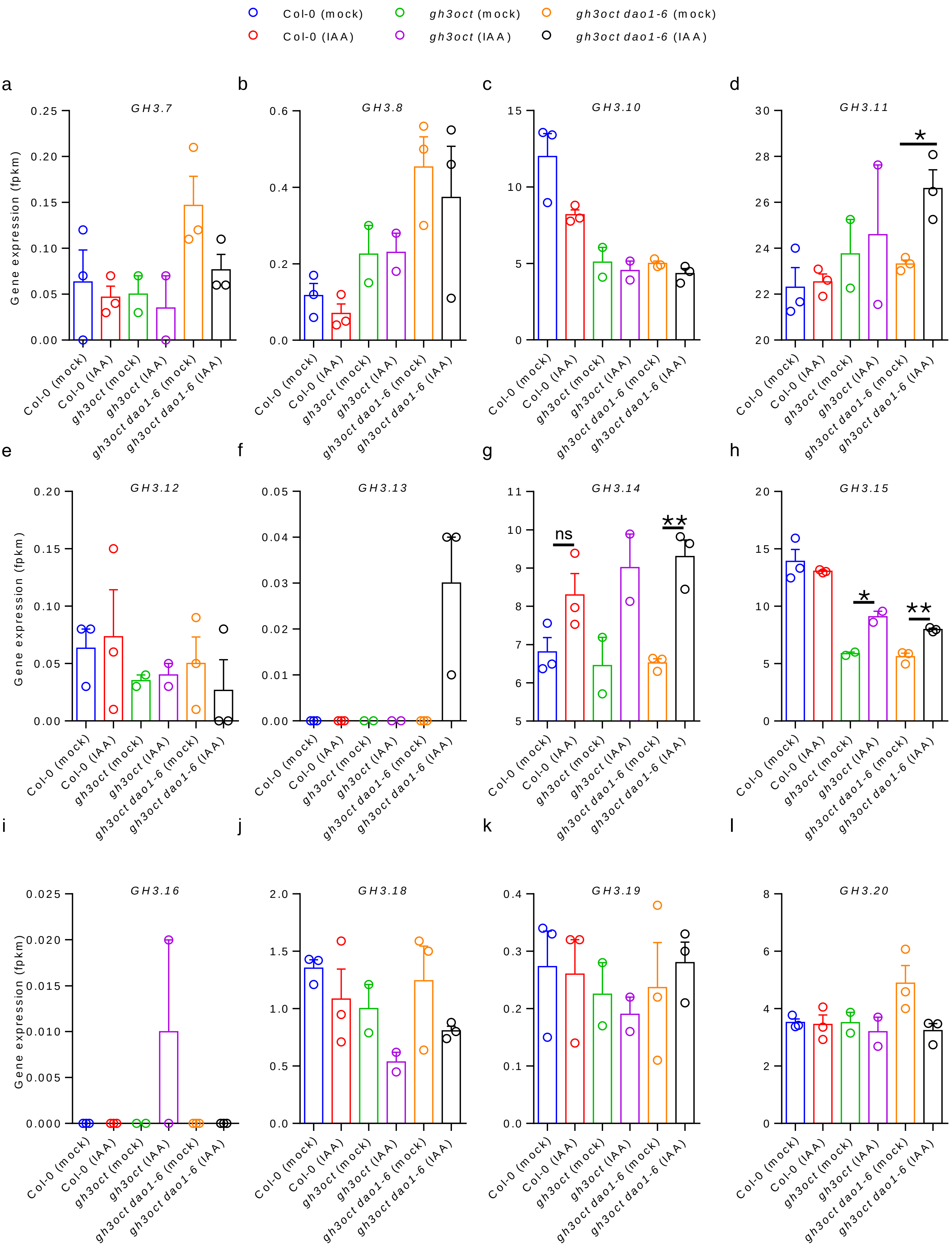


**Figure S12**. Differences in the expression of non-group II *GH3* genes. Expression levels are reported as fragments per kilobase of transcript per million reads mapped (FPKM). Error bars indicate the standard error of the mean. For each genotype, it is represented by mock (left) and IAA (right) treatments. Asterisks indicate values that are significantly different from the mock value in a Student´s *t*-test (*p<0.05, **p<0.01).

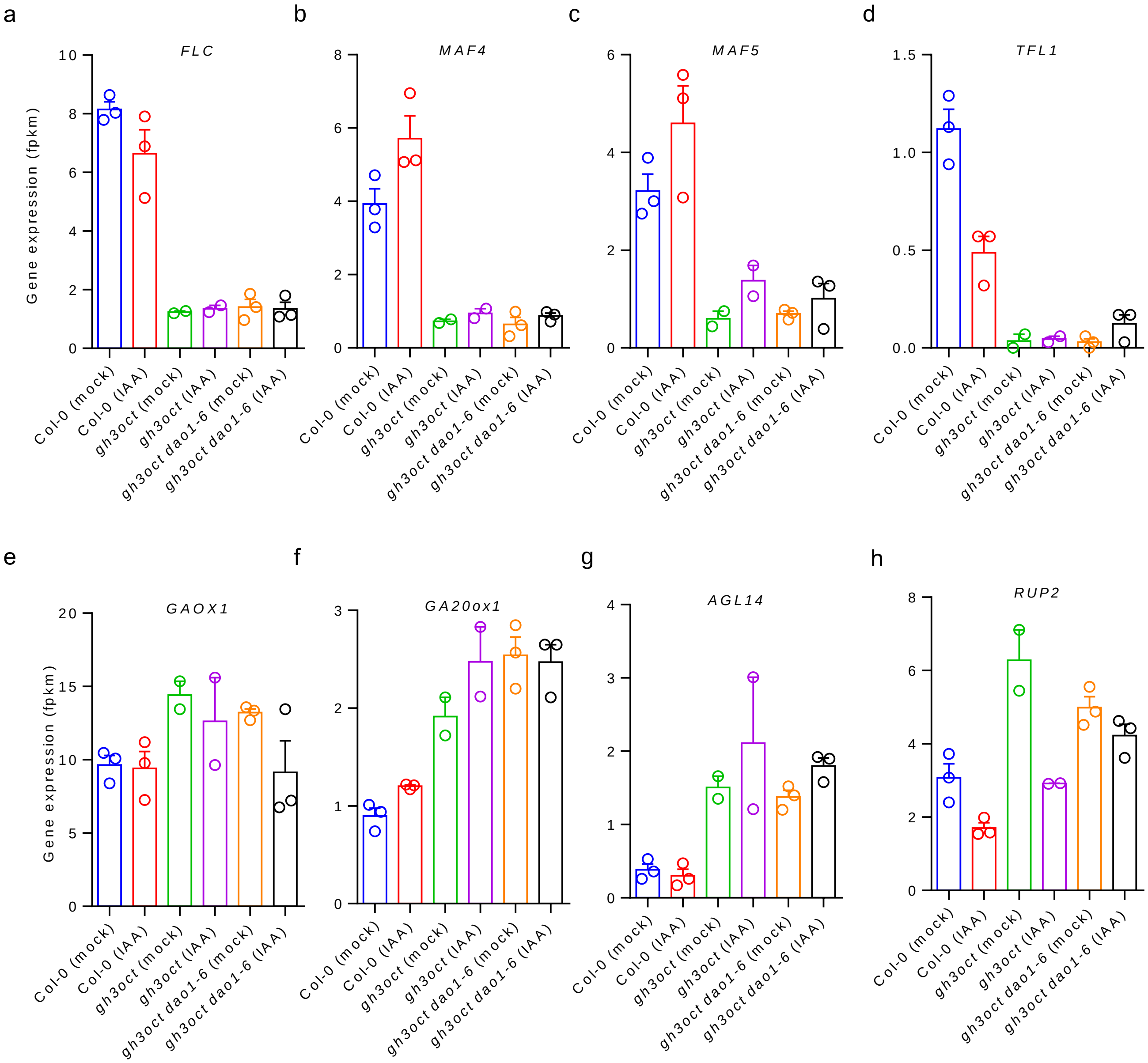


**Figure S13**. Differences in the expression of flowering time-related genes. Expression levels of flowering (a-d) repressor and (e-h) activator genes, expressed in fragments per kilobase of transcript per million reads mapped (fpkm). Error bars indicate standard error of the mean. For each genotype, it is represented by mock (left) and IAA (right) treatments.
