## Supplementary Tables for "Comprehensive characterisation of IAA inactivation pathways reveals the impact of glycosylation on auxin metabolism and plant development"

**Table S1. Primer sets used in this work**

| Purpose | Primer name | Primer sequence (5' → 3') | |
|---|---|---|---|
| | | Forward sequence (F) | Reverse sequence (R) |
| CRISPR/ Cas9 cloning | DAO1_g#1 | AATGTCTCTTTCAACGTCCATACG | AAACCGTATGGACGTTGAAAGAGA |
| | DAO1_g#2 | gTACGCACTTTCACCTCGTAgttttagagctatgctg | TACGAGGTGAAAGTGCGTAcaatcactacttcgactc |
| | DAO1_g#3 | GAGCTCTGTAACCACTCCCTgttttagagctatgctg | AGGGAGTGGTTACAGAGCTCaatcactacttcgactc |
| | DAO1_g#4 | gACGCTTGCTATCAACCTCGgttttagagctatgctg | CGAGGTTGATAGCAAGCGTcaatcactacttcgactc |
| | DAO1_g#5 | gGTACCCCTTCGTGACTAATgttttagagctatgctg | ATTAGTCACGAAGGGGTACcaatcactacttcgactc |
| | DAO2_g#6 | gCCGATGTGTTACTAGGGAAgttttagagctatgctg | TTCCCTAGTAACACATCGGcaatcactacttcgactc |
| | DAO2_g#7 | gTTAGCGGAGAGCTACGGAGgttttagagctatgctg | CTCCGTAGCTCTCCGCTAAcaatcactacttcgactc |
| Genotyping | UBQ10pro | GTGATCAAGGTAAATTTCTGTGTT | TGAGAAATTGAAATCTGAATTGTG |
| | B-module | ATGTGAGTTAGCTCACTCATTAG | CAACTGTTGGGAAGGGCGAT |
| | G-10332 | CTCGTATGTTGTGTGGAATTGTGAGC | |
| | G-37561 | TCAGACCTAGAAAAGCTGCAAA | |
| | G-23253 | CCAAACGTAAAACGGCTTGT | |
| | G-40025 | TTGGTCTTGAACAAGAGATC | |
| | 1.2en1.1p-F | AAGCTGATTTGGTTCTATTGAACTA | |
| | G-33037 | GTGTGCGCAATGAAACTGATGC | |
| | G-37561 | TCAGACCTAGAAAAGCTGCAAA | |
| | mCherry-R1 | | GGTGCCGCGCAGCTTCACCT |
| | mCherry-R2 | | CCTCGCCCTTGCTCACCAT |
| | Cas9 | ATGGATAAGAAGTACTCTATCGG | AACCTTCCTCTTCTTCTTAGGAT |
| | dao1-5 dao2-2 (wt 3021 / mut 461) | CGGAGTCATCATTCCGACTA | GGCTACGgtgggatgatatc |
| RT-qPCR | qGH3.1 | GGACAACTCGGTTGGACCAT | TGCACCTCTTGAGATTGCGT |
| | qGH3.2 | TGCGTGAGCTTCACACCTAT | CTAAAACCGCACATCATCCG |
| | qGH3.3 | CTCCGTGCCATTGGATTCCT | ATCAGCCAGTTCTTGGTCCG |
| | qGH3.4 | TGATCGGTGTGAGGCTTACG | TGGAGATACGTGTGGTGCAG |
| | qGH3.5 | TGCTCCAATTATCGAGCTATTGA | TGTTTGTGACCAGGAACCCA |
| | qGH3.6 | ACCTATGCTGGGCTTTACAGG | GCGGCATATGAAGCTGAACTG |
| | qGH3.9 | CGACGATGAACAAGTCCCCT | GGTAACGGTACAATCCTGCGA |
| | qGH3.17 | CATTTGTTAAGTTGCTAATTGGTGT | AGAGGACTTTGCTGAAAGTTTGT |
| | qACT2 | CCGCTCTTTCTTTCCAAGC | CCGGTACCATTGTCACACAC |

| Sample | Q20 % | Clean read number | SRA reference |
|---|---|---|---|
| Col-0(Mock)_1 | 97.66 | 87,755,182 | SAMN16288319 |
| Col-0(Mock)_2 | 97.53 | 85,625,344 | SAMN16288320 |
| Col-0(Mock)_3 | 97.47 | 85,531,692 | SAMN16288321 |
| Col-0(IAA)_1 | 97.61 | 87,287,732 | SAMN16288322 |
| Col-0(IAA)_2 | 97.73 | 87,578,238 | SAMN16288323 |
| Col-0(IAA)_3 | 97.69 | 83,689,142 | SAMN16288324 |
| gh3oct(Mock)_1 | 97.38 | 87,368,762 | SAMN16288325 |
| gh3oct(Mock)_2 | 97.52 | 83,285,346 | SAMN16288326 |
| gh3oct(IAA)_1 | 97.72 | 87,589,734 | SAMN16288327 |
| gh3oct(IAA)_2 | 97.94 | 87,211,028 | SAMN16288328 |
| gh3octdao1(Mock)_1 | 97.74 | 87,755,182 | SAMN16288329 |
| gh3octdao1(Mock)_2 | 97.51 | 86,567,062 | SAMN16288330 |
| gh3octdao1(Mock)_3 | 97.43 | 87,126,212 | SAMN16288331 |
| gh3octdao1(IAA)_1 | 97.44 | 87,145,640 | SAMN16288332 |
| gh3octdao1(IAA)_2 | 98.40 | 84,716,878 | SAMN16288333 |
| gh3octdao1(IAA)_3 | 98.00 | 81,573,998 | SAMN16288334 |

**Supplementary Table 2**. Sequencing statistics of the different RNA-seq samples. All the experiments were performed using paired-end (PE) reads of 100 bp in length. Q20 % values indicate the percentage of reads (forward, left; reverse, right) that reach a Phread score of 20 or higher.

**Table S3**. Modules used for the GreenGate-based cloning of the CRISPR-Cas9 plasmids generated in this work.

| Construct | Supermodule | Modules | Insert |
|---|---|---|---|
| #1 | M | A | EC1.2enhancer–EC1.1promoter |
| | | B | *A.thaliana* codon-optimized Cas9 |
| | | C | rbcS terminator |
| | | D | DAO2_g#7 |
| | | E | DAO1_g#4 |
| | | FH | F-H adapter (pGGG001) |
| | N | HA | H-A adapter (pGGG002) |
| | | A | UBQ10 promoter |
| | | B | mCherry CDS |
| | | C | rbcS terminator |
| | | D | DAO2_g#6 |
| | | E | DAO1_g#3 |
| | | F | HygR (pGGF005) |
| #2 | - | A | EC1.2enhancer–EC1.1promoter |
| | | B | *A.thaliana* codon-optimized Cas9 |
| | | C | rbcS terminator |
| | | D | DAO1_g#2 |
| | | E | DAO1_g#5 |
| | | F | HygR (pGGF005) |
